# Fast computational screening of extrusion parameters for bioprinting of skeletal myoblasts in alginate hydrogel

**DOI:** 10.64898/2026.10.19.760174

**Authors:** Zahed Tavangari, Ali Maghami, Antonio Papangelo, Giada Graziana Genchi

## Abstract

**Background:** Real-time selection of extrusion bioprinting parameters requires rapid prediction of an operating window that balances material throughput against mechanically induced cell damage. Full numerical simulations can resolve the relevant flow mechanics, but their computational cost limits repeated evaluation during process adjustment. In this work, a finite element–Gaussian process (GP) framework is developed for rapid operating-window prediction in skeletal myoblast bioprinting for tissue engineering purposes.

**Methods:** An axisymmetric model of inertia-retained, shear-thinning flow of a 1.5% (w/v) alginate suspension of L8 rat myoblasts is coupled to a previously documented damage law that separates nozzle shear from entrance extensional injury of the same cell model. The dataset comprises 32 combinations of four nozzle diameters (100–330 µm) and eight applied pressures (25–400 kPa). GP surrogates are developed to predict flow rate, wall shear stress and total mechanical damage.

**Results:** GP surrogates respond in under one millisecond *per* parameter combination, compared with approximately 1.5 hours *per* finite element case. In leave-one-out validation, the surrogates reproduce the simulated damage with a mean absolute error of 0.34 percentage points and the flow rate with a mean absolute percentage error of 6.8%. With a whole nozzle diameter withheld, the damage error rises to 0.88 percentage points. With screening limits of predicted damage below 20% and a flow rate of at least 1 µL/s, 17 of the 32 simulated conditions lie within the operating window. The damage decomposition also shows the dependency of nozzle selection on the imposed constraint: at 200 kPa, the predicted damage increases from 9.0% to 18.0% with nozzle diameter increasing from 200 µm to 330 µm. The 330 µm nozzle carries a 15.8-fold higher flow rate than the 200 µm nozzle, which raises the effective extensional stress at the nozzle entrance. At a matched flow rate of 1 µL/s, the predicted damage instead decreases from 5.5% to 4.6%.

**Conclusions:** The developed framework supports real-time computational screening of printing parameters within the sampled domain and suggests the need for careful selection of nozzle diameter and applied pressure together, since the ordering of nozzles by predicted damage reverses between fixed pressure and matched flow rate.

## Background

Bioprinting enables the controlled deposition of cell-laden materials to produce tissue constructs with complex 3D architecture, including aligned skeletal muscle with the potential to fill large anatomical gaps and exhibit contractile activity for motor function recovery [1–3]. In recent decades, the formulation of muscle-specific bioinks and the development of sophisticated extrusion systems have supported the fabrication of skeletal muscle constructs with different degrees of success [4,5]. Obtaining functional constructs, however, requires a careful and timely choice of those printing conditions that can preserve cell viability while sustaining accurate material deposition. Real-time decisions in extrusion bioprinting can be taken by the implementation of models for the rapid identification of the printing conditions satisfying both throughput targets and limits on the predicted mechanical damage. During extrusion, changes in the ink properties or delivery conditions can significantly alter the flow obtained at a certain pressure value. For this reason, real-time flow measurements and automated pressure adjustments have been implemented to improve dispensing consistency and compensate for ink inhomogeneities [6]. However, a predictive model can significantly complement this real-time feedback by proactively evaluating the mechanical consequences of candidate printing parameters prior to extrusion [7,8].

In this context, the main biological constraint is represented by the mechanical loading imposed on cells as the bioink passes through the cartridge and nozzle. On one hand, the mechanical loading is governed by extrusion pressure or flow rate, nozzle diameter and length, nozzle contraction geometry, and bioink rheology [9–11]. On the other hand, cell damage depends on shear stress, extensional stress, exposure time, and cell-specific susceptibility to damage. In particular, substantial extensional injury at the cartridge–nozzle interface has been shown to affect cell viability in addition to shear stress along the nozzle [9,12,13]. Nozzle design studies have also examined how contraction angle and outlet diameter affect shear and extensional loading and post-printing viability [14]. Needle-mediated injection experiments have additionally identified shear rate as an important predictor of cell injury under the tested conditions [15]. Overall, the experimental evidence in the literature highlights the need to interpret damage laws based on the cell types and rheological conditions for which they were established [15]. Pressure and nozzle diameter must therefore be considered together when defining a predicted mechanical-damage operating window.

Finite element, finite volume, and lattice-Boltzmann methods provide numerical descriptions of the flow and cell deformation during extrusion, while damage formulations connect these mechanical fields to biological outcomes [7,16,17]. Relevant applications of these methods include the work of Ning *et al*., who separated shear and extensional injury with cell-specific experiments [9], and of Han *et al*., who developed a pressure-work-based damage model [18]. Interestingly, Chirianni *et al*. combined axisymmetric finite element modeling (FEM), a generalized law accounting for shear and extensional damage, and reduced-order modeling to construct pressure–diameter nomograms for quick inspection of flow rate and cell viability prior to printing [8]. Nevertheless, repeated high-fidelity evaluations remain to date computationally demanding. At fixed element order, nested-dissection direct factorization on classical two-dimensional regular meshes has an operation count 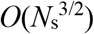, where *N*_s_ is the number of spatial degrees of freedom [19]. Repeating such a factorization over *I* nonlinear iterations gives the illustrative scaling 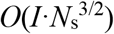, and the actual cost depends on the solver, mesh, and convergence behavior. Geometry and material or loading parameters influence the spatial refinement of the model and its convergence: in the present study, we show that the numerical cost remains parameter-dependent through the mesh, nonlinear convergence, and trajectory integration, motivating a surrogate for repeated operating-window evaluation. When entrance loading is estimated as the difference between applied pressure and developed nozzle friction, upstream losses and small changes in the friction estimate can also materially affect the inferred extensional contribution. Assessing this sensitivity is therefore important when interpreting damage predictions.

Machine learning (ML) and data-driven methods offer a way to move this computational effort to an offline stage. In mechanics, neural networks have predicted mechanical properties in real time [20], while neural operators have learned families of fluid-flow solutions [21]. In bioprinting, Reina-Romo *et al*. combined computational fluid dynamics with Gaussian process analysis to investigate nozzle geometry and shear stress [7]. They also demonstrated that increasing outlet radius can increase maximum shear stress at fixed pressure, while reducing it at fixed flow rate. Thus, both surrogate-assisted nozzle analysis and the dependence of hydrodynamic trends on the imposed constraint are established precedents. Blanco *et al*. reviewed computational nozzle optimization and proposed a design-of-experiments framework for selecting bioink-specific operating conditions [22]. Other data-driven models have linked elastic modulus and yield stress to printability [23], and the importance of ML-assisted formulation and process optimization has been noted by Freeman *et al*. [24]. Generating training data and fitting a model can be expensive, but a trained neural surrogate with a fixed computational graph and input/output sizes requires a fixed operation count *per* query, independent of the numerical values of the process parameters [25]. This permits approximately uniform latency across the explored domain on fixed hardware. Millisecond inference is feasible with suitable implementations and hardware. For instance, Li *et al*. reported 5 ms for a Fourier neural operator on a 256 × 256 flow grid [21]. GP regression is particularly suited to small simulation datasets and also provides posterior uncertainty [26]. This uncertainty describes the surrogate approximation conditional on its training data and assumptions.

Building on previous surrogate-assisted nozzle studies and reduced-order process-design tools [7,8], we propose an L8 cell-specific computational framework for rapid evaluation of operating conditions under simultaneous throughput and mechanical-damage constraints (schematized in Fig. 1). We investigate whether narrower nozzles remain the more damaging once the damage law includes entrance extension, and whether the damage ordering, like the shear-stress ordering, depends on holding the pressure or the flow rate fixed. We quantify how entrance extensional loading, nozzle shear exposure, and inertia influence predicted damage and can affect nozzle selection under fixed-pressure and matched-throughput conditions. We also assess the sensitivity of the inferred entrance contribution to upstream friction and the pressure-drop estimation method. A GP surrogate makes the resulting pressure–diameter response available for repeated evaluation without additional flow simulations. Our original contribution lies in the quantitative connection between these mechanisms and the selected operating conditions within the investigated bioink and geometry. Our analysis uses a single 32-case dataset, with particle tracking providing supporting information on the radial distribution of nozzle shear exposure. The resulting predicted mechanical-damage operating window is intended for screening within the sampled domain.

**Fig. 1.**
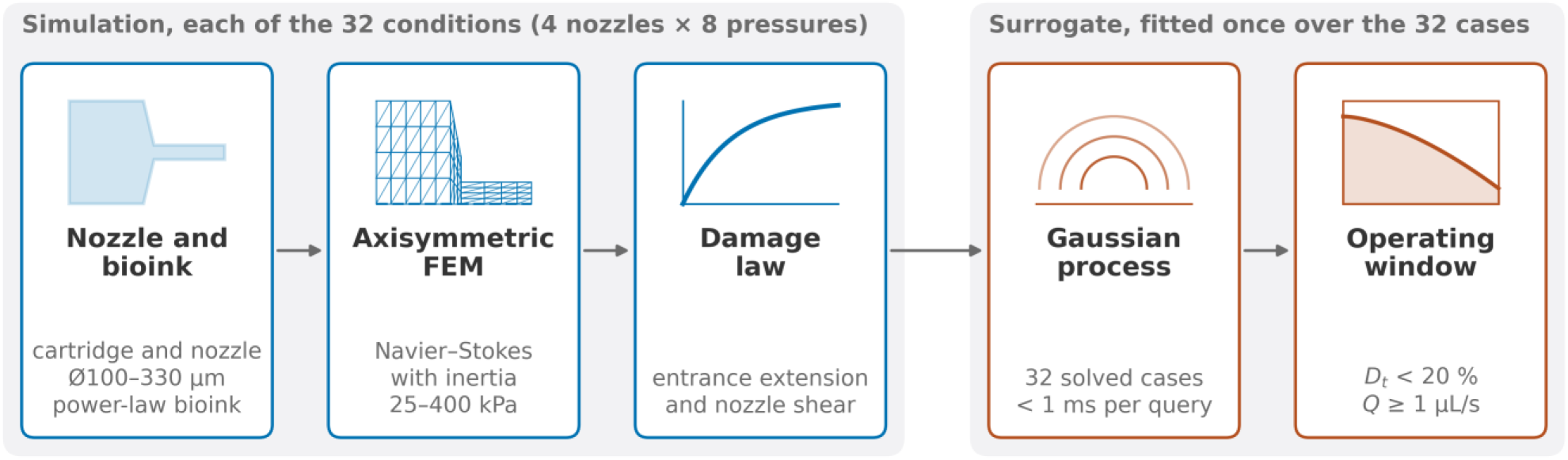
Study workflow, from nozzle geometry to operating window. For each of the considered 32 combinations of nozzle diameter and pressure, the axisymmetric finite element model computes the flow, from which the damage law gives the entrance and nozzle-shear damage. A Gaussian process fitted to the 32 cases gives the operating window. Lagrangian tracers are integrated at the reference condition only (200 kPa, 200 µm diameter nozzle).

## Methods

### Nozzle geometry and rheological assumptions

The computational domain reproduces the cartridge and nozzle assembly published elsewhere [9], with a 4.2 mm cartridge bore contracting to a 2.0 mm cartridge outlet and interchangeable nozzles of 100, 200, 250 and 330 µm diameter (Fig. 2). The cartridge is a 10.5 mm long cylinder, with a 1.5 mm long truncated-cone transition for nozzle anchoring, and the nozzle is a 10 mm long cylinder, for a total flow path of 22.0 mm. The bioink is modeled as an incompressible shear-thinning fluid with the power-law parameters that Ning *et al*. derived from measured mass flow rates for a 1.5% (w/v) alginate suspension of L8 myoblasts with 1 × 10^6^ cells/mL.

**Fig. 2.**
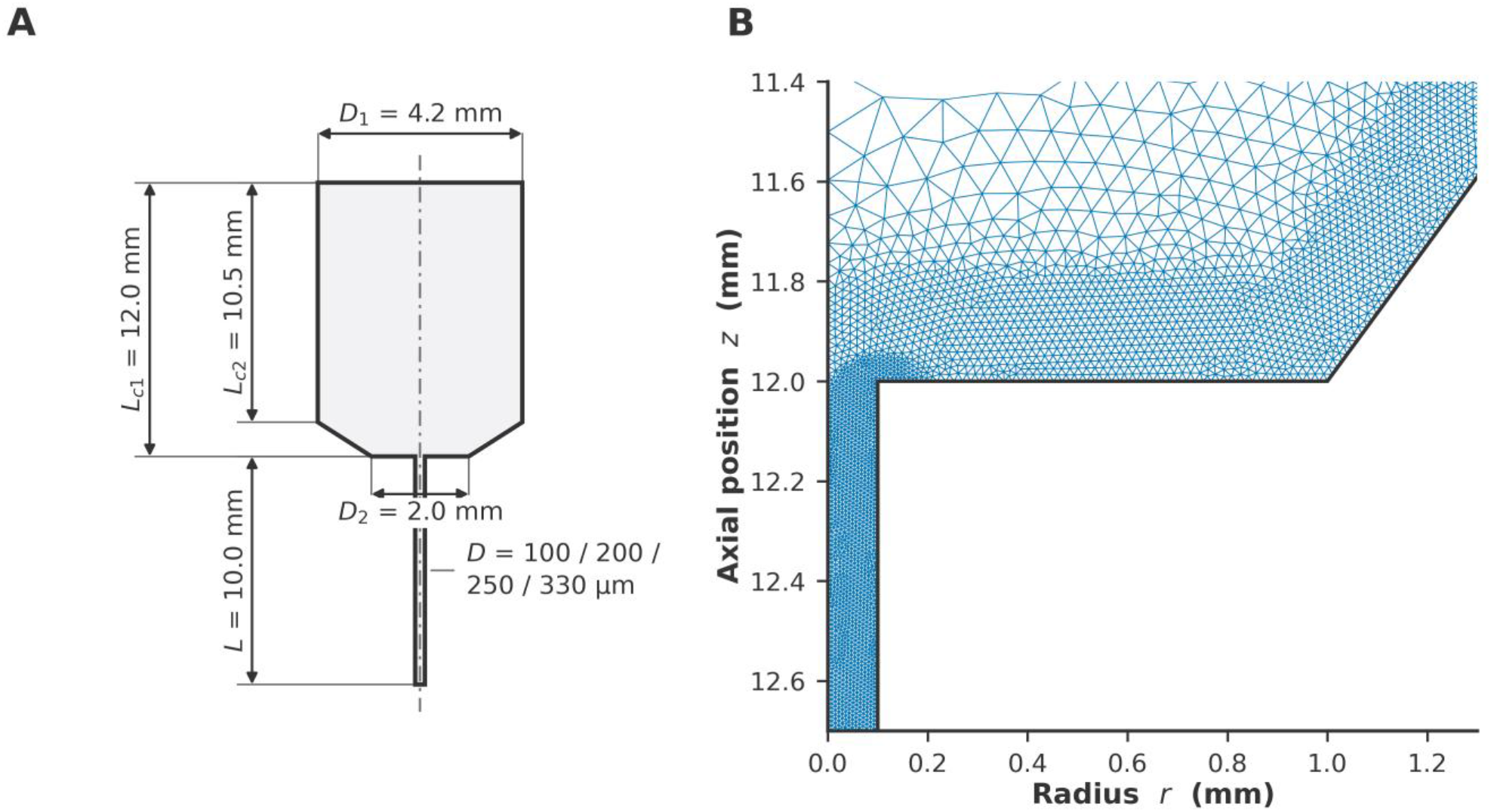
Cartridge and nozzle geometry from [9]. (A) Axisymmetric domain (from top to bottom): cartridge bore D_1_ = 4.2 mm, cartridge straight length = 10.5 mm, truncated-cone transition length = 1.5 mm (half-angle 36.3°), transition outlet D_2_ = 2.0 mm, nozzle diameter D = 100, 200, 250 or 330 µm and length L = 10.0 mm. L_c1_ = 12.0 mm includes the transition. Radial axis exaggerated 2.13×. (B) The contraction region is depicted in true aspect ratio, with the mesh.

### Axisymmetric flow model

The cell-laden alginate suspension is modeled as an incompressible, generalized-Newtonian fluid undergoing steady, laminar, axisymmetric flow. These assumptions are consistent with the flow model used in [9]. The governing equations are written in the axisymmetric generalized-Newtonian form adopted in previous work [8]. Cylindrical coordinates (*r, θ, z*) are used, with the axial coordinate *z* aligned with the cartridge and nozzle. The velocity field is assumed to have no circumferential component and no dependence on *θ*,

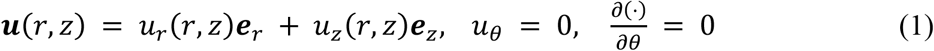

where ***e***_*r*_ and ***e***_*z*_ are the radial and axial unit vectors, and ∇_axi_ denotes the gradient and divergence in axisymmetric cylindrical coordinates. Gravity and other body forces are neglected, since the hydrostatic pressure over the 22.0 mm flow path, 0.21 kPa, is 0.85% of the lowest applied pressure, 25 kPa. Steady flow, with convective inertia retained, then obeys conservation of volume and the momentum balance:

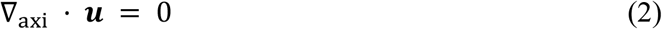

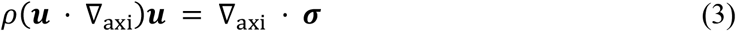

where *ρ* is the suspension density and the Cauchy stress tensor is

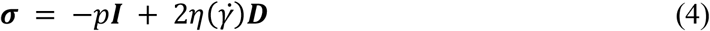

Here, *p* is the pressure, ***I*** the identity tensor and *η* the apparent viscosity, given below as a function of the shear-rate magnitude 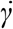, and

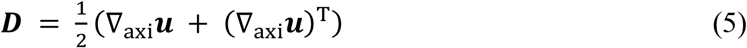

is the rate-of-deformation tensor. For an axisymmetric flow without swirl, its nonzero components are

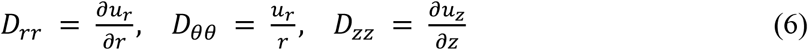

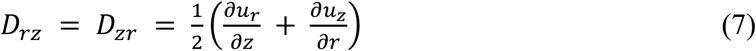

Accordingly, the axisymmetric incompressibility condition is

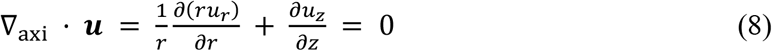

and the generalized shear-rate magnitude is defined by the second invariant of ***D***,

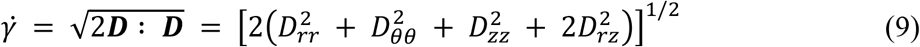

Following [9], the yield stress τ_0_ of the published flow law is set to zero, as they did for this low alginate concentration when fitting their power-law parameters, and the apparent viscosity is described by the power-law relation:

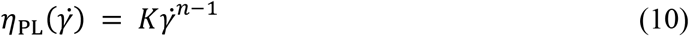

with consistency index *K* = 25.60 Pa·s^n^, flow-behavior index *n* = 0.35, and density *ρ* = 990 kg m^−3^. The boundary of the meridional domain is decomposed as

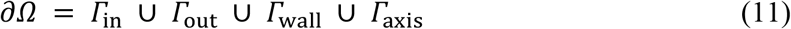

where *Γ*_in_ is the cartridge inlet, *Γ*_out_ the nozzle outlet, *Γ*_wall_ the solid walls and *Γ*_axis_ the symmetry axis. The applied pressure ΔP is imposed as a normal-traction condition. By using the outward unit normal ***n***, the boundary conditions are

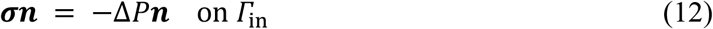

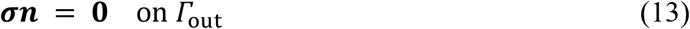

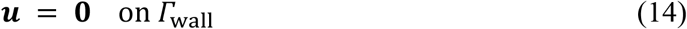

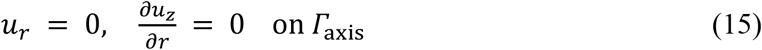

Eq. 12 applies the prescribed pressure difference as a compressive inlet traction. Eq. 13 is a zero-total-traction, or “do-nothing,” outlet condition. It is not a strong Dirichlet condition on the pressure variable. For a fully developed outflow, for which the normal viscous stress vanishes, this condition reduces to zero gauge pressure at the outlet.

Steady generalized-Newtonian flow is solved with DOLFINx 0.10 [27] by using Taylor– Hood elements (P_2_ velocity, P_1_ pressure) on anisotropic Gmsh meshes refined toward the nozzle, with twelve elements across the nozzle radius (mesh M12). Because the power-law viscosity diverges at zero shear rate, it is smoothly bounded between an upper limit of 500 Pa·s and a lower limit of 0.003 Pa·s; the lower limit lies above the high-shear viscosity of alginate inks in [8]. The upper bound acts only where the shear stress is below 5.2 Pa; a developed-flow estimate puts its flow-rate effect below 0.3%. The nonlinear system is solved by Newton iteration, each step with the MUMPS direct solver through PETSc; Additional file 1 gives the regularization of the viscosity and of the axisymmetric terms (Section S1, Eqs. S1–S7) and the finite element weak form (Section S2, Eq. S8). Continuation is used in the lower viscosity bound, from 10 to 0.003 Pa·s in five levels, in the applied pressure and in density. A case is kept only if the solve converges at full density, and all 32 do.

Eight applied pressures (25, 50, 75, 100, 150, 200, 300 and 400 kPa) are solved for each of the four nozzle diameters, at 1.5 hours of compute *per* case on average (0.9 to 2.3 hours; Additional file 1: Section S1). The simulations run in a DOLFINx 0.10 container under HTCondor on the ReCaS-Bari cluster. All 32 cases are also solved on a finer mesh, M16, with sixteen elements across the nozzle radius, and on M12 without the inertial terms. Between the two meshes, the predicted total damage changes by at most 0.033 percentage points (Additional file 1: Fig. S2A) and the flow rate by at most 0.22%. The increase in predicted total damage from the 200 to the 330 µm diameter nozzle at 200 kPa is 101.0% on M12 and 100.9% on M16.

### Comparison with published flow and stress data

The model is compared, with no calibration factor, with the 200 µm diameter data from [9] over 50 to 400 kPa, that is the measured mass flow rates and the computed wall shear stresses. The wall shear stress, formed from the solved flow rate, agrees with the values computed by Ning *et al*. to within 0.47% between 50 and 400 kPa for the 200 µm diameter nozzle, and to within 0.54% across all seven conditions of their computation (Table 1). The measured flow rate is over-predicted, by 4.5% at 200 kPa and by 19.7% to 36.7% at the other pressure values, within 0.1 to 2.0 standard deviations of the measurements.

**Table 1.** Wall shear stress and entrance pressure: present model compared with previously published data [9].

| D<br>($\mu\text{m}$) | $\Delta P$<br>(kPa) | $\tau_w$, model<br>(Pa) | $\tau_w$, Ning<br>(Pa) | $\Delta\tau_w$<br>(%) | $p_{\text{en}}$, model<br>(kPa) | $p_{\text{en}}$, Ning<br>(kPa) | $\Delta p_{\text{en}}$<br>(%) | Re |
| --- | --- | --- | --- | --- | --- | --- | --- | --- |
| 100 | 200 | 495.8 | 496 | −0.04 | 1.674 | 1.650 | +1.45 | 0.012 |
| 200 | 50 | 245.7 | 246 | −0.14 | 0.867 | 0.867 | +0.05 | 0.002 |
| 200 | 100 | 491.3 | 491 | +0.06 | 1.741 | 1.773 | −1.79 | 0.047 |
| 200 | 200 | 981.2 | 981 | +0.02 | 3.756 | 3.734 | +0.60 | 1.222 |
| 200 | 300 | 1459.8 | not reported | n/a | 8.038 | not reported | n/a | 7.949 |
| 200 | 400 | 1900.0 | 1909 | −0.47 | 20.000 | 18.246 | +9.61 | 27.535 |
| 250 | 200 | 1213.3 | 1214 | −0.05 | 5.865 | 5.620 | +4.37 | 5.194 |
| 330 | 200 | 1523.8 | 1532 | −0.54 | 15.301 | 14.345 | +6.66 | 26.488 |

D, nozzle diameter; $\Delta P$ , applied pressure; $\tau_w$ , wall shear stress from the solved flow rate, Eq. 16; $p_{\text{en}}$ , entrance pressure, Eq. 17; Ning, the wall shear stresses and contraction pressure drops computed by Ning *et al.* for L8 cells [9]; $\Delta$ , difference from their value; Re, Reynolds number.

The wall shear stress τ_w_ is formed from the solved volumetric flow rate *Q* through the Rabinowitsch wall shear rate γ? _w_ for a power-law fluid in a nozzle of radius R, which is exact for developed flow in a straight tube [9]:

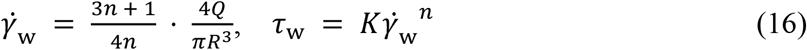

The entrance pressure p_en_ is the residual of the applied pressure ΔP after the pressure drop Δp_tip_ of developed flow along the nozzle length L_tip_ = 10 mm:

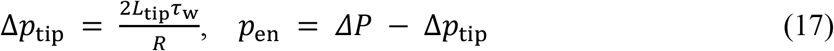

The Reynolds number of each case is Re = ρ·2Rū/η_w_, where ū = Q/(πR^2^) is the mean velocity in the nozzle and η_w_ = K(ū/R)^n−1^ is the viscosity at the characteristic shear rate ū/R. The finite element wall stress serves only as a check on the analytic τ_w_, which it matches to within 0.61% at the reference condition.

The reference condition used throughout is 200 kPa through the 200 µm diameter nozzle. There the solved flow rate is 17.94 µL/s, and Eqs. 16 and 17 give a wall shear stress of 981 Pa and an entrance pressure of 3.76 kPa.

The full residual of Eq. 17 is used because Ning *et al*. identify the entrance pressure with the Bagley end correction, which leaves upstream friction in the intercept, and computed it from Navier–Stokes simulations that retain inertia. They fitted the extensional branch of their damage law to such entrance pressures for the 200 µm diameter nozzle between 50 kPa and 400 kPa. At 200 kPa, the present residual lies within 0.6 to 6.7% of their entrance pressures across the four diameters, and within 1.8% between 50 and 200 kPa for the 200 µm diameter nozzle, where the difference reaches 9.6% at 400 kPa (Table 1). For a 200 µm diameter nozzle, the residual lies 1.7% above a Bagley extrapolation on the present model (Additional file 1: Fig. S2C).

### Mechanobiological damage model

Cell damage is quantified by using the Ning *et al*. formulation calibrated to L8 myoblasts [9], in which total damage is decomposed into shear-driven and extensional components combined through a cascade rule. The Ning *et al*. model parameters used in this study are given in Table 2.

**Table 2.** Parameters of the damage law for L8 myoblasts from [9], as used here.

| Parameter | Symbol | Value | Component |
| --- | --- | --- | --- |
| Shear pre-factor | $a_1$ | $3.20 \times 10^{-4}$ | Shear ($D_{st}$) |
| Shear stress exponent | $b_1$ | 0.61 | Shear ($D_{st}$) |
| Residence time exponent | $c_1$ | 0.52 | Shear ($D_{st}$) |
| Extensional pre-factor | $a_2$ | $1.43 \times 10^{-3}$ | Extensional ($D_{et}$) |
| Extensional stress exponent | $b_2$ | 0.55 | Extensional ($D_{et}$) |

Shear damage is evaluated on the analytic capillary profile [18], on which the local shear stress τ_s_ varies linearly with the radial coordinate r:

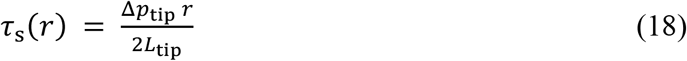

and the axial velocity of developed power-law flow, v_z_(r), distinct from the solved axial velocity u_z_ of Eq. 1, is

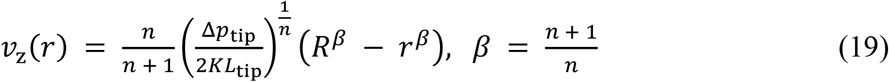

A cell entering the nozzle at radius r thus spends a residence time t_s_(r) = L_tip_/v_z_(r) in the nozzle and accumulates the shear damage D_s_(r), with a_1_, b_1_ and c_1_ from Table 2:

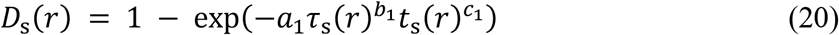

Averaged over the nozzle cross-section, following the convention of Ning *et al*. [9], D_s_(r) gives the shear damage of the nozzle, D_st_:

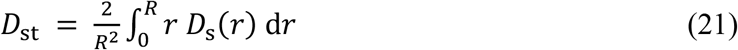

The flow-weighted average, the damage of the cells collected at the outlet, is 0.182% for the 200 µm diameter nozzle at 200 kPa, compared with 0.286% for the area average (Additional file 1: Eq. S9 and Table S1).

In the extensional branch, which follows Cogswell’s converging-flow analysis [28], the entrance pressure p_en_ of Eq. 17 is converted into an effective extensional stress τ_e_:

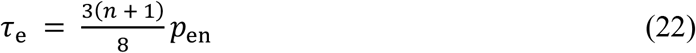

which causes the extensional damage D_et_, with a_2_ and b_2_ from Table 2:

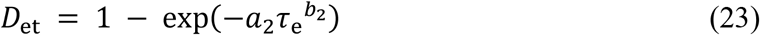

The cascade rule of Ning *et al*. [9], according to which a cell must survive the entrance before it is exposed to the wall, combines the two branches into the total damage D_t_:

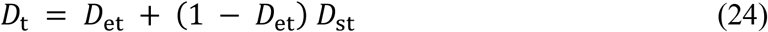

Cogswell’s converging-flow analysis relates the entrance pressure to an effective extensional stress [28], and the same pressure corresponds to the capillary end correction characterized by Bagley [29]. τ_e_ is therefore an effective stress derived from the entrance pressure under the Cogswell– Ning closure, not a resolved extensional stress. The residual also carries the wall friction of the cartridge and the truncated-cone transition and an inertial part, both quantified in the Results.

Ning *et al*. calibrated the shear branch between 100 and 1000 Pa for 10 to 120 s and obtained the extensional branch from L8 myoblast printing runs with extensional stresses spanning 440 to 9251 Pa [9]. The reference condition and the fixed-pressure diameter comparison both lie inside the calibrated extensional range, at τ_e_ = 1902 Pa and 7746 Pa. Across the grid, eight of the 32 cases fall below 440 Pa and five exceed 9251 Pa; these extrapolations are marked † in Table 3 and Additional file 1: Table S1.

**Table 3.**
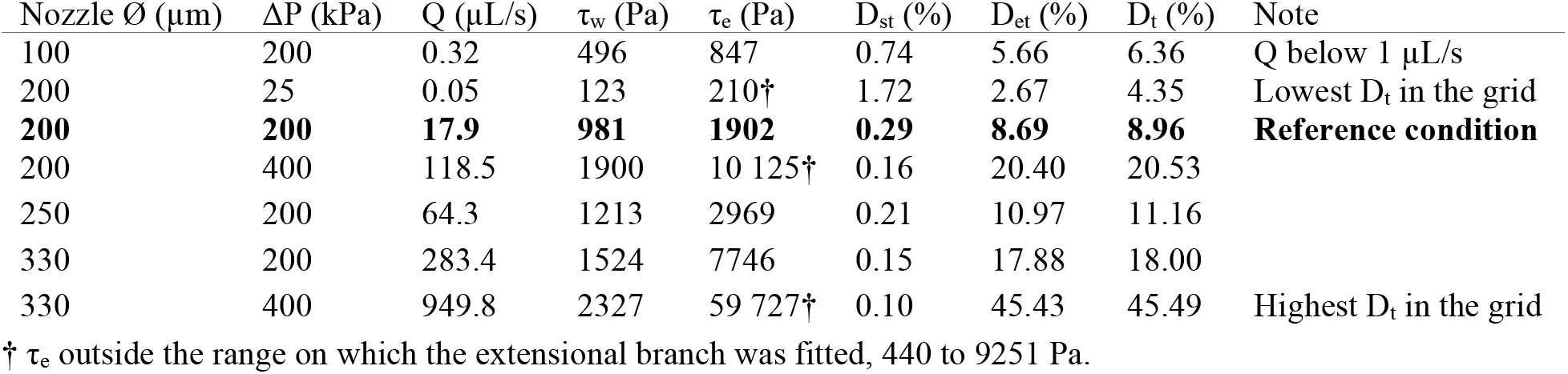
Flow, stress and damage from the finite element model for seven of the 32 diameter–pressure combinations.

The shear law as used here reproduces the twenty shear-damage points in the supplementary data of Ning *et al*. with R^2^ = 0.906. The wall stress exceeds the 1000 Pa ceiling of its calibration in nine of the 32 cases, where D_st_ reaches at most 0.21 percentage points. It lies below the 100 Pa floor only for the 100 µm diameter nozzle at 25 kPa, at τ_w_ = 62 Pa. Nozzle residence times also lie below the 10 to 120 s of the shear calibration in 30 of the 32 cases, so D_st_ is extrapolated in exposure time as well (Additional file 1: Section S3 and Table S1).

### Lagrangian particle tracking

To resolve the exposure of individual cells, 168 tracer particles are seeded uniformly by area over the inner 95% of the cartridge radius, on a release plane 5 mm into the cartridge. The seeding radius is r_i_ = 0.95 R_in_ √u_i_, with u_i_ uniform on (0, 1) and R_in_ = 2.1 mm. Each tracer is advected through the solved velocity field with the explicit Runge–Kutta 5(4) integrator (RK45) of SciPy to the nozzle outlet, and trajectories that touched a wall are discarded. Fig. 3C tracers are seeded by the same rule 0.2 mm from the inlet.

**Fig. 3.**
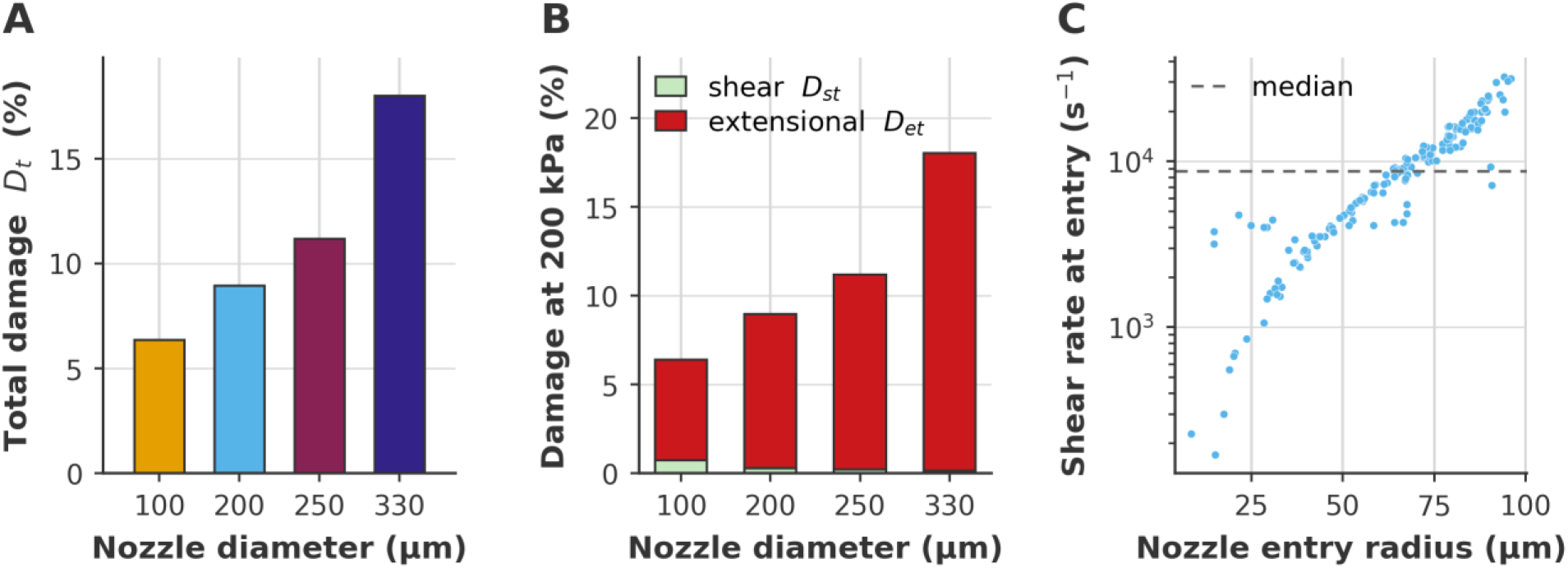
Damage of the finite element solutions against nozzle diameter at 200 kPa. (A) Total damage. (B) The same cases split into entrance and nozzle shear terms. (C) Entry shear rate against entry radius, 200 µm diameter nozzle, for 168 tracers released 0.2 mm from the inlet; dashed line, median 8698 s^−1^.

Each tracer takes the shear damage of a radial profile of the straight nozzle at the radius at which it enters the nozzle, which is exact wherever the flow is fully developed. The profile is formed from the mean shear stress and axial velocity in 160 radial bins of the solved field, with residence time t_s_(r) = L_tip_ / v_z_(r). It is scaled so that its area average reproduces the analytic D_st_ of Eq. 21. D_et_, derived from the bulk entrance pressure, is uniform across the tracers of a case, and the cascade rule gives the *per*-particle total damage.

### Gaussian process surrogate

GP regression suits the small simulation budget, 32 cases at about 1.5 hours each in a two-dimensional input space, because it interpolates without a fixed functional form and reports a posterior uncertainty [26]. Three GPs are fitted independently, for log_10_ *Q*, log_10_ τ_w_ and D_t_, with applied pressure and nozzle diameter as inputs. Each uses a Matérn 5/2 kernel with separate length scales for pressure and diameter, multiplied by a constant variance, with hyperparameters set by maximizing the marginal likelihood from ten random starts (Additional file 1: Section S4). The diagonal term α = 10^−6^ is a fixed regularization in standardized target units, not a measured noise. In the Results, FEM values are finite element solutions passed through the damage law, of which the 32 grid cases train the surrogate, and surrogate values are GP predictions.

The surrogate is scored by leaving out one case at a time, a whole nozzle diameter or a whole applied pressure, with the input standardization and hyperparameters refitted inside every fold. A missing interior diameter or pressure tests interpolation, and a missing edge tests extrapolation beyond the remaining levels. For each output the mean absolute, root-mean-square and largest errors are reported, with the fraction of held-out values inside the nominal 95% posterior interval. The conditions misclassified under the two operating-window criteria, D_t_ < 20% and *Q* ≥ 1 µL/s, are counted as well. Linear, quadratic and cubic polynomial surfaces are scored on the same splits.

### Operating-window definition

The operating window is defined by two simultaneous criteria, a predicted damage D_t_ < 20% and a throughput (volumetric flow rate) *Q* ≥ 1 µL/s at the nozzle outlet. The damage limit is a screening limit on predicted mechanical damage, since the previously published data for L8 myoblasts show no pressure threshold. Proliferation of printed cells relative to control falls with applied pressure, from 86.7% at 50 kPa to 71.7% at 400 kPa 72 h after printing [9]. The throughput limit is a practical rate for depositing millimeter-scale constructs in a standard experimental session.

For each nozzle diameter, the surrogate is evaluated at 3751 applied pressures evenly spaced over the trained range (25–400 kPa), and the lowest predicted D_t_ satisfying both criteria is recorded as the operating point. Eleven further conditions are then fixed from the surrogate predictions before any of them is solved. They are the operating point at each of the four solved diameters and at two unsolved ones, 150 and 290 µm, and the pressure at which the predicted D_t_ reaches 20% wherever it lies in the trained range. The eleventh is an interior condition at 150 µm and 300 kPa. They are solved with the same model and settings as the grid, and none is used to fit the surrogate.

### Data analysis and visualization

The GP surrogate and the analysis are run in Python 3.13 with NumPy 2.2, SciPy 1.15 and scikit-learn 1.7. All figures are drawn with Matplotlib 3.10 from the simulation output, and the animation in Additional file 2 is rendered with PyVista from the solved velocity field, with the particle paths integrated in SciPy. The code and the results behind every figure are available in the repository named under Availability of data and materials, where each figure is regenerated by its own script.

### Use of generative artificial intelligence

Claude (Anthropic) and GitHub Copilot (GitHub) were used for language editing and assistance with code development and debugging. All outputs were reviewed and verified by the authors, who take full responsibility for the content of the article. These tools were not used to generate or alter data, results, tables, figures, or scientific conclusions.

## Results

### Predicted damage across nozzle diameter and applied pressure

Because the shear-dominated view predicts that a wider nozzle reduces damage [1,30], the predicted damage is first compared across nozzle diameters at fixed applied pressure. Across the 32 conditions, the predicted total damage D_t_ ranges from 4.35% to 45.49% and depends on both nozzle diameter and applied pressure (Table 3). At fixed pressure, D_t_ increases with nozzle diameter (Fig. 3A), because a wider nozzle carries a much higher flow rate, which raises the entrance velocity and the effective extensional stress (Fig. 4A). At ΔP = 200 kPa, D_t_ is 8.96, 11.16 and 18.00% when nozzles with a diameter of 200 µm, 250 µm and 330 µm are considered, implying a 9.0-percentage-point increase within the considered nozzle diameter range.

**Fig. 4.**
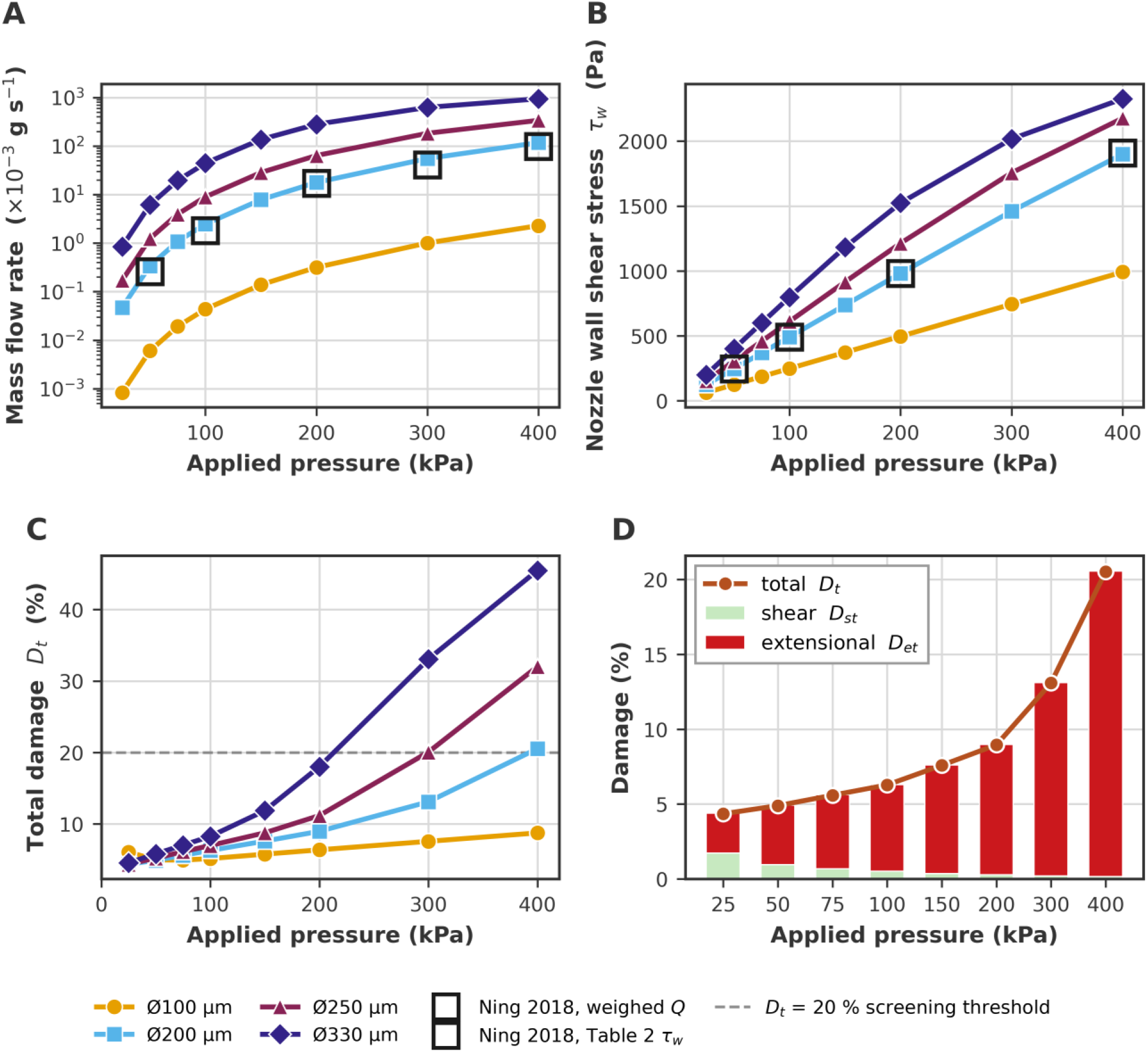
Flow rate, wall shear stress and damage of the 32 finite element solutions against applied pressure. (A) Mass flow rate, with black square frames for the flow rates weighed by Ning et al. (B) Nozzle wall shear stress from the solved flow rate (Rabinowitsch relation), against the values computed by Ning et al. (Table 1). (C) Total damage D_t_, with a dashed line at the 20% damage limit. (D) The damage split into D_et_ and D_st_ for the 200 µm diameter nozzle.

The predicted total damage increase is monotonic in diameter at every pressure of 75 kPa or more (Fig. 4C). At 25 and 50 kPa, the 200 µm diameter nozzle gives the lowest D_t_, by only 0.004 percentage points over the 250 µm diameter nozzle at 25 kPa. The 100 µm diameter nozzle is the most damaging one, with D_t_ = 6.05% at 25 kPa, compared with D_t_ = 4.35% for the 200 µm diameter nozzle. The slow transit through the narrow nozzle gives a shear contribution of 4.34% and an entrance contribution of only 1.80%.

D_et_ contributes 89.0–99.3% of D_t_ across the four diameters at 200 kPa (Fig. 3B), and 29.7% at the low-pressure corner of the grid (100 µm, 25 kPa; Additional file 1: Fig. S2B). D_st_ decreases from 0.29% for a 200 µm diameter nozzle to 0.15% for a 330 µm diameter nozzle (Table 3), because the shorter residence time at the higher flow rate reduces D_st_ more than the higher wall shear stress raises it (Fig. 4B). Across the 168 tracers that enter the 200 µm diameter nozzle at 200 kPa, the shear rate at nozzle entry rises toward the wall, from 171 to 32 244 s^−1^, with a median of 8698 s^−1^ (Fig. 3C).

At a pressure of 200 kPa, the flow rate shows a 15.8-fold increase from *Q* = 17.94 µL/s for a 200 µm diameter nozzle to *Q* = 283.4 µL/s for a 330 µm diameter nozzle. At fixed applied pressure, a wider nozzle thus delivers more material and produces more predicted damage.

At the reference condition (ΔP = 200 kPa, 200 µm diameter nozzle), D_et_ = 8.69%, D_st_ = 0.29% and D_t_ = 8.96%, so extensional damage accounts for 97.1% of the predicted total. Of the 3.756 kPa entrance pressure, 0.474 kPa, or 12.6%, is fully developed wall friction in the cartridge and the truncated-cone transition, and 6.6% is inertial, from the solution without inertia. The inertial share exceeds 50% in six of the 32 cases, all at high flow rate (Additional file 1: Table S1).

Shear damage stays small because the nozzle residence time is short and shear thinning limits the wall shear stress to 981 Pa, which does not strongly activate the shear term of the Ning *et al*. formulation. The extensional term is the larger of the two at every pressure for the three wider nozzles. Conversely, for the 100 µm diameter nozzle, the damage changes from shear-dominated to extensional-dominated between 25 and 50 kPa.

Raising the applied pressure increases D_et_ at every diameter, because a higher entrance velocity drives a larger p_en_ and τ_e_, and reduces D_st_, because cells pass through the nozzle faster (Fig. 4D, for the 200 µm diameter nozzle). With D_et_ dominant, D_t_ increases with pressure at every diameter except for the 100 µm diameter nozzle, where the shear-dominated low-pressure corner produces an interior minimum near 75 kPa that a shear-only analysis would not predict.

Between 200 and 400 kPa, D_t_ rises from 8.96 to 20.53% for a 200 µm diameter nozzle, and from 18.00 to 45.49% for a 330 µm diameter nozzle, where the extensional term is largest (Fig. 4C).

Tracking of 168 particles to the nozzle outlet allows calculation of nozzle-entry radius for each particle, along with shear damage (Fig. 5A–C). Near-wall particles experience both the longest residence time and the highest shear stress. The former increases from 11.5 ms at the nozzle axis to 135 ms at 97.6% of the nozzle radius, 97.6 µm from the axis of the 200 µm diameter nozzle and 2.4 µm from its wall, where the outermost tracer enters the nozzle. *Per*-particle shear damage therefore varies from 0.04% on the axis to 0.66% for that tracer. Because the extensional damage is the same for every particle (8.69%), the particle-to-particle differences in total damage come only from the shear term: across the 168 particles, the total damage varies by 0.564 percentage points, 6.3% of its mean value (Fig. 5D). The velocity field through the contraction is shown in Additional file 2.

**Fig. 5.**
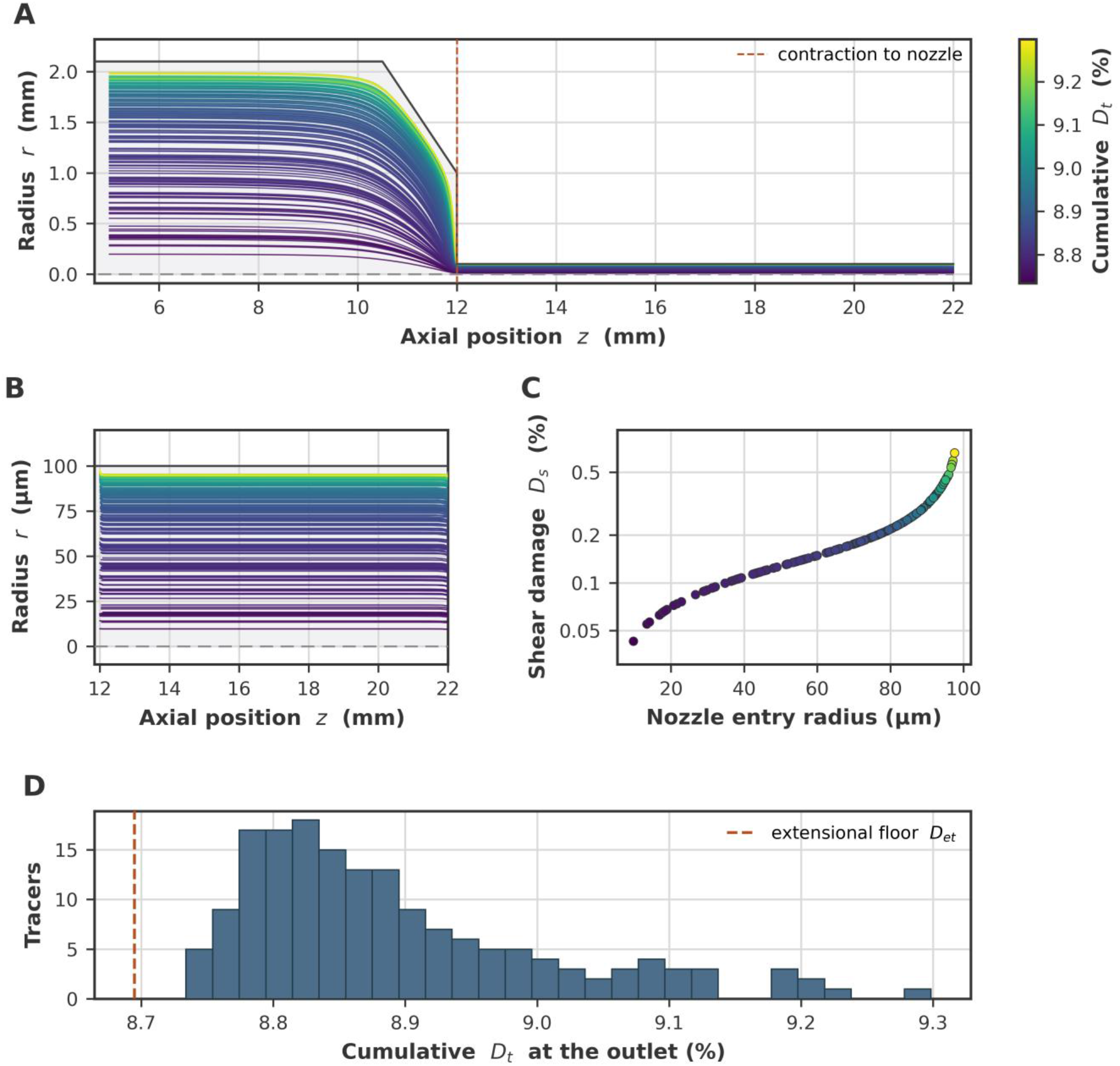
Lagrangian trajectories through the cartridge and nozzle. The 168 tracers are seeded uniformly by area and integrated through the finite element velocity field to the outlet. (A) Tracks colored by the damage each tracer carries; axial and radial coordinates use independent scales. (B) Tracks inside the nozzle. (C) Shear damage (with logarithmic axis) against entry radius. (D) Distribution over the tracers of D_t_ above a common extensional floor of 8.69 percentage points.

### GP surrogate accuracy and operating window

Over the pressure–diameter plane, the surrogate gives response surfaces of flow rate, wall shear stress and total damage (Fig. 6A–C) and the posterior standard deviation of the damage (Fig. 6D). With each case left out in turn, the surrogate reproduces the 32 solved values of D_t_ with a mean absolute error of 0.338 percentage points, a root-mean-square error of 0.683 and a largest error of 2.48. This error is 2.75 times smaller than that of the best polynomial surface, the cubic at 0.931 percentage points (Additional file 1: Table S3). In flow rate, the mean absolute error is 0.027 in log_10_ *Q*, a factor of 1.064. Computed directly from the held-out predictions, the mean absolute percentage error in *Q* is 6.8%, with a largest error of 78%. Applied to the two operating-window criteria, the held-out predictions place the same 17 conditions inside the window and the same 15 outside it as the finite element results, so no condition is misclassified.

**Fig. 6.**
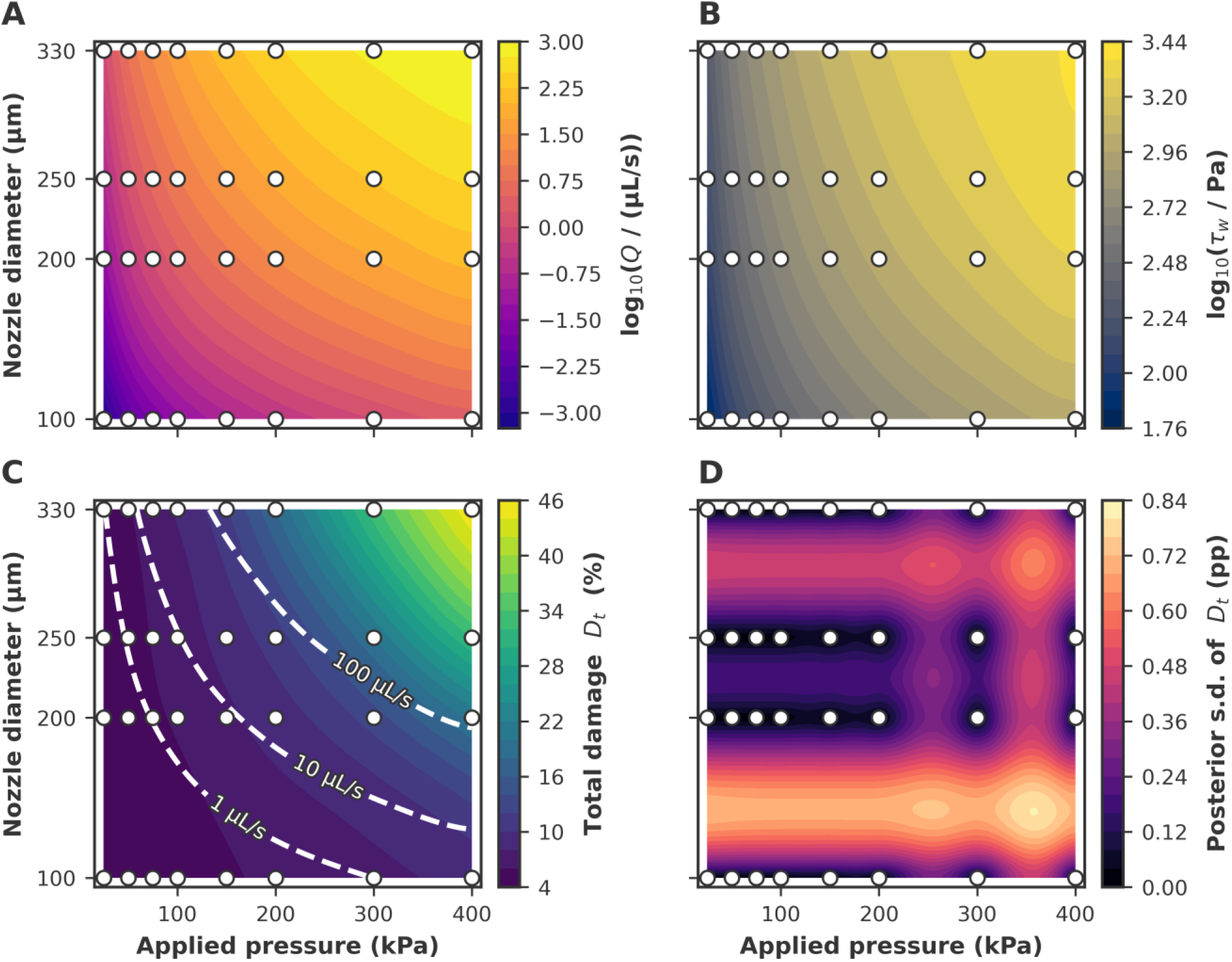
Gaussian process surrogate response surfaces over applied pressure and nozzle diameter. (A) log_10_ Q, Q in µL/s. (B) log_10_ τ_w_, τ_w_ in Pa. (C) Total damage D_t_ (%), with iso-flow contours at 1, 10 and 100 µL/s. (D) Conditional GP posterior standard deviation of D_t_ (percentage points). Markers are the 32 FEM training cases.

With a whole diameter held out, the error in D_t_ rises to 0.883 percentage points on average. It is 0.543 at the two interior diameters, where the surrogate interpolates, and 1.223 at the 100 and 330 µm diameter edges, where it extrapolates (Additional file 1: Fig. S1 and Table S2). The flow rate is more sensitive to a missing diameter, with errors of 0.060 in log_10_ *Q*, a factor of 1.15, at the interior diameters and 0.278, a factor of 1.90, at the edges. With a whole pressure held out, the error in D_t_ is 0.194 percentage points at interior pressures and 1.498 at the edges. Holding out a diameter places one in-window condition outside the window and no out-of-window condition inside it, while holding out a pressure places two out-of-window conditions inside the window and one in-window condition outside it.

With each case left out in turn, the nominal 95% interval of the GP posterior contains the finite element D_t_ in 27 of 32 cases (84.4%; Additional file 1: Fig. S3). With a whole diameter held out, the intervals on log_10_ *Q* contain none of the held-out values. The posterior standard deviation is therefore used here only to map where the surface is least constrained by the training cases (Fig. 6D).

Of the 32 solved conditions, 17 satisfy both criteria (Fig. 7A). At 200 kPa, all four nozzles stay below the 20% damage limit, but only the three wider ones reach 1 µL/s, whereas the 100 µm diameter nozzle delivers 0.32 µL/s. At 400 kPa, the predicted total damage reaches 20.53% for the 200 µm diameter nozzle and 45.49% for the 330 µm diameter nozzle.

**Fig. 7.**
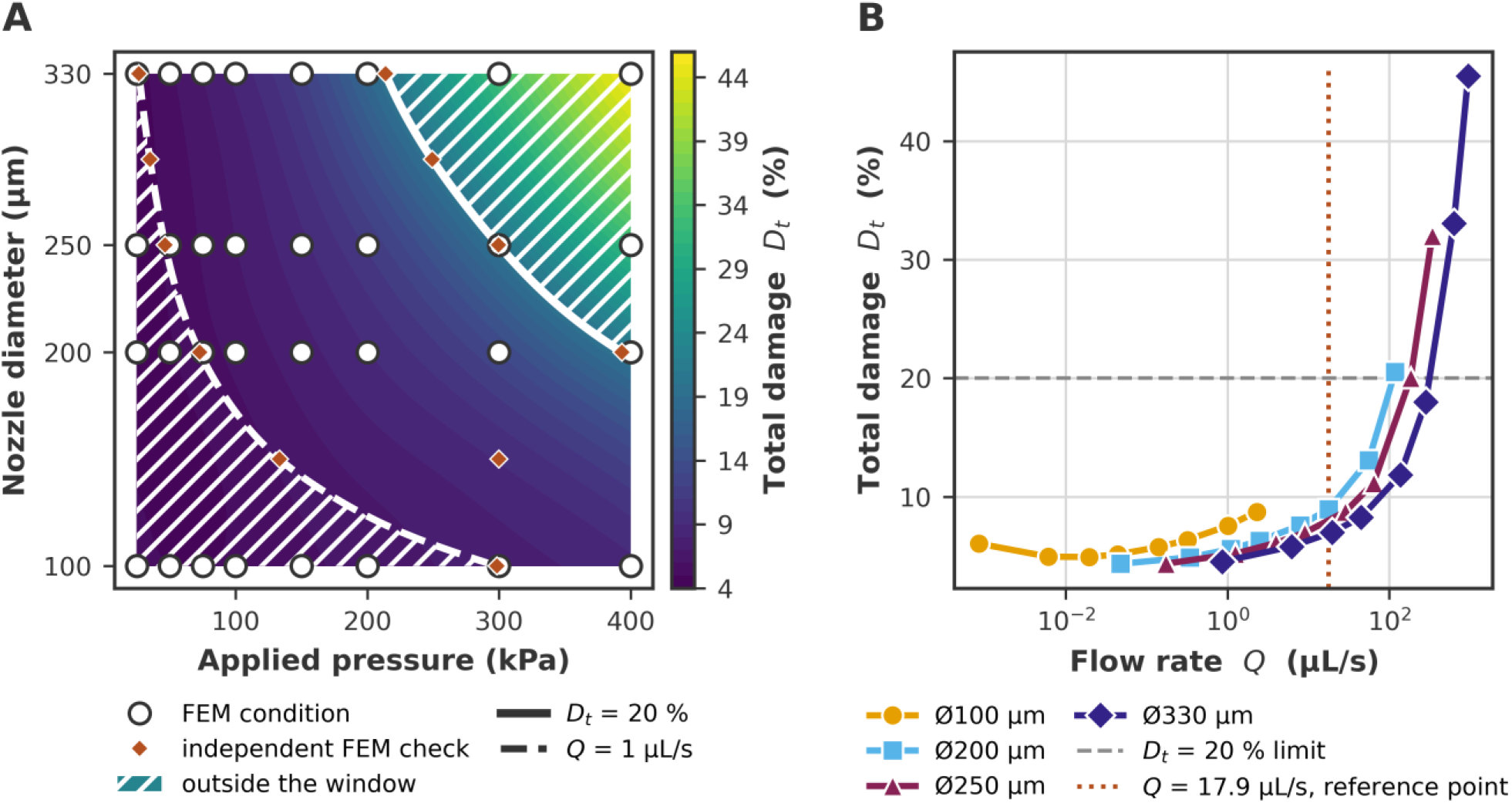
Operating window and the effect of the applied constraint. (A) Total damage from the GP surrogate, with the two criteria, D_t_ = 20% and Q = 1 µL/s, and circles for the 32 FEM conditions, 17 of which satisfy both. Diamonds are the eleven FEM checks, fixed from the surrogate before solving and not used to train it (Additional file 1: Table S5). (B) Total damage of the 32 FEM conditions against volumetric flow rate for each nozzle.

The surrogate places the operating point of every diameter on the flow limit. The operating pressures are 298.5 kPa for 100 µm, 72.7 kPa for 200 µm, 46.5 kPa for 250 µm and 26.6 kPa for 330 µm diameter nozzles, with predicted D_t_ of 7.54, 5.52, 5.08 and 4.61%. Above the operating pressures, the predicted damage reaches the 20% limit, which bounds the window from above, at 393.4 kPa for 200 µm, 299.6 kPa for 250 µm and 213.8 kPa for 330 µm diameter nozzles. For the 100 µm diameter nozzle, D_t_ stays below 20% up to 400 kPa, the end of the trained range. The eleven FEM checks, solved after the surrogate had fixed them, reproduce its damage predictions to within 0.40 percentage points, 0.068 on average, and its flow rate to within 5.7% (Fig. 7A; Additional file 1: Table S5). The largest errors fall at the unsolved diameters, in D_t_ at 150 µm and in flow rate at 150 µm and 290 µm.

The 95% interval of the damage prediction contains the FEM value in all eleven checks, and that of the flow rate in eight. The window decision differs in three checks, each placed on a limit by design. Two lie on the damage limit, where the FEM gives D_t_ = 19.95% and the surrogate 20.01%, and one on the flow limit, where the FEM gives *Q* = 0.9998 µL/s and the surrogate 1.0026 µL/s. With the 17 in-window grid conditions, these checks test the continuous window between the solved cases.

Refitted with a Matérn 3/2 kernel, or with α = 10^−4^, the surrogate moves none of the operating pressures by more than 0.3 kPa, and with α = 10^−2^ it moves them by up to 7.4 kPa (Additional file 1: Table S4). The rougher kernel places 30 of the 32 single held-out cases inside its nominal 95% interval, but its error with a diameter held out is 2.7 times larger.

### Contribution of inertia to the diameter effect

To separate the viscous and inertial parts of the diameter effect, the whole 32-case grid was re-solved on the same domain and mesh without the inertial terms and post-processed with Eqs. 16–24. Between 200 and 330 µm diameter at 200 kPa, the predicted damage rises by 101.0% with inertia retained and by 32.7% without it, so about two thirds of the fixed-pressure diameter effect is inertial.

Inertia accounts for 0.0, 3.4, 12.6 and 36.2% of the predicted damage for 100, 200, 250 and 330 µm diameter nozzles at 200 kPa, and its share of the damage reaches 53.1% at the widest nozzle over the full pressure range. The Reynolds number is 1.2 for 200 µm and 26.5 for 330 µm diameter nozzles at 200 kPa, rising to 194.8 for the 330 µm diameter nozzle at 400 kPa. An inertial contribution of this magnitude is therefore expected.

The inertial part of p_en_ matches the kinetic energy the jet carries out of the nozzle, 1.56 ρū^2^/2, where 1.56 is the kinetic-energy factor of the developed power-law profile. The two agree to within 7% in the twelve cases where the inertial part exceeds 0.1 kPa, so this part is the pressure spent accelerating the ink. Cogswell’s relation, reported in Eq. 22, was derived for creeping flow, and the inertial part enters it only through the calibration: Ning *et al*. fitted a_2_ and b_2_ to entrance pressures from Navier–Stokes solutions that include it. Their calibration conditions, 200 µm diameter nozzle at pressures from 50 to 400 kPa, carry inertial shares of 0 to 58.9% in the present model. The fixed-pressure comparison for the 330 µm diameter nozzle at 200 kPa, amounting to 59.1%, lies at that edge. The four further cases with larger shares, for the 250 and 330 µm diameter nozzles at 300 or 400 kPa, also lie outside the calibrated τ_e_ (Additional file 1: Table S1).

### Diameter ordering at matched flow rate

With volumetric throughput held fixed instead of applied pressure, the same solved cases give the opposite ordering. Interpolated at a matched flow rate of 1 µL/s, monotonically in log_10_ *Q*, D_t_ is 7.53% for 100 µm, 5.53% for 200 µm, 5.08% for 250 µm and 4.59% for 330 µm diameter nozzles. These values lie within 0.02 percentage points of the surrogate operating points, 7.54, 5.52, 5.08 and 4.61% (Fig. 7A). The widest nozzle now gives the lowest predicted damage, because a wider nozzle reaches a given throughput at a lower entrance velocity and the extensional damage rises with that velocity. The inversion also holds at *Q* = 17.94 µL/s, the flow rate of the reference condition (Fig. 7B), where D_t_ is 8.96% for 200 µm, 7.90% for 250 µm and 6.90% for 330 µm diameter nozzles. The 100 µm diameter nozzle does not reach that flow rate in the grid.

### Flow at the cartridge-to-nozzle contraction

At the reference condition, the flow converges from the 1.0 mm radius of the cartridge outlet into the 100 µm radius of the nozzle over a short axial distance (Fig. 8A). Along the centerline, the velocity rises from 0.005 to 0.873 m s^−1^, and the extensional strain rate peaks at 5.44 × 10^3^ s^−1^ over a full width at half maximum of about 130 µm (Fig. 8B).

**Fig. 8.**
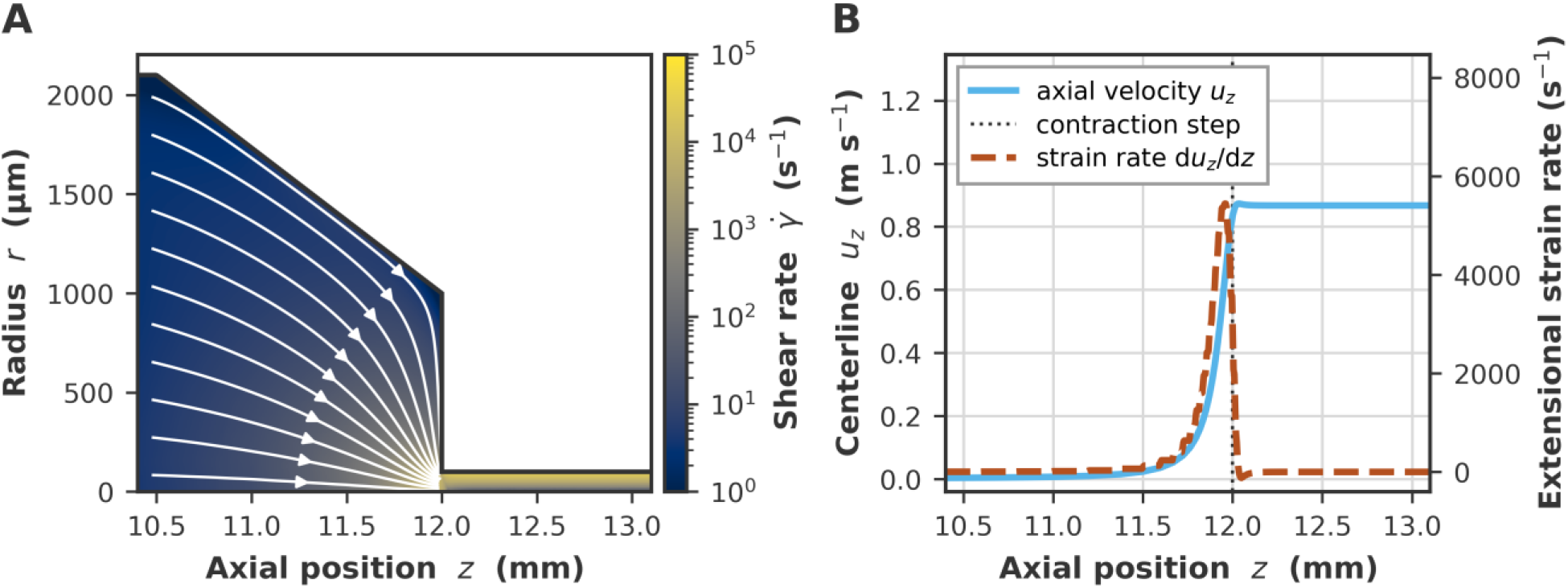
Finite element flow field at the cartridge-to-nozzle contraction. 200 µm diameter nozzle at ΔP = 200 kPa, considered as the reference condition. (A) Shear rate with streamlines integrated to the step; the radial axis spans the 2.1 mm cartridge radius, so the truncated-cone transition and the 10:1 step are at true proportions. (B) Centerline velocity and the implied extensional strain rate.

## Discussion

### Nozzle diameter at fixed pressure and at matched flow rate

In the proposed model, a wider nozzle raises the predicted damage at fixed applied pressure (by 101.0% from the 200 to the 330 µm diameter nozzle at 200 kPa), and lowers it at matched flow rate (Figs. 3 and 7B). Both directions follow from the decomposition. The diameter response is carried almost entirely by the entrance term, which rises with entrance velocity. At fixed pressure, a wider nozzle raises that velocity; at matched flow rate, it lowers it.

At fixed applied pressure, the damage Ning *et al*. measured in the same L8 myoblast cell line increased with nozzle diameter from 100 to 330 µm, as predicted here, although not significantly [9]. Through the same law, their extensional stresses give 8.7% for 200 µm and 17.3% for 330 µm diameter nozzles, close to the present 8.69% and 17.88%, implying similar entrance loading in the two models. At a constant flow rate of 3 × 10^−3^ g/s (corresponding to ~3.0 µL/s), Ning *et al*. report the opposite ordering in RSC96 Schwann cells: 16.84% damage through a 100 µm diameter nozzle and 7.28% through a 330 µm diameter one [9]. The present model predicts both orderings for L8 myoblasts on one grid, and the dependence on the constraint is consistent with the joint action of applied pressure and nozzle diameter on post-printing survival [11,31].

The direction of the diameter effect is unchanged by mesh refinement and by removing inertia, while its magnitude depends on inertia, which carries about two thirds of the fixed-pressure increase, and on how the nozzle friction is computed. At the reference condition, the solved wall shear is 975.2 Pa and the analytic value 981.2 Pa, implying a difference of 0.61%. Because the residual is the small difference of two large pressures, the 0.61% difference moves the entrance pressure from 3.76 to 4.96 kPa, and D_et_ from 8.69 to 10.06%.

The rise of damage with diameter at fixed pressure runs against the common expectation in extrusion bioprinting that smaller nozzles are more damaging [1,30], which is an expectation grounded in shear-dominated analyses [32]. That expectation holds at matched flow rate, where a wider nozzle lowers the wall shear stress (Eq. 16), but at fixed applied pressure the wall shear stress itself rises with nozzle diameter (Fig. 4B). Measured damage was also found to rise with nozzle diameter at fixed pressure in human dermal fibroblasts printed in alginate through 200 to 400 µm diameter needles [18] and, although not significantly, in RSC96 Schwann cells [9].

In the present geometry, the step from the 2.0 mm cartridge outlet into the nozzle is steep, with a diameter ratio of 6.1 when considering the 330 µm diameter and of 20 when considering the 100 µm diameter. As a result, the entrance stress exceeds the nozzle wall stress throughout the parameter space, by a factor of at least 1.59. Selecting the widest available nozzle to reduce damage at a fixed applied pressure would increase it on these predictions.

For process design, the flow rate required by the print should therefore be fixed first, and nozzle diameter and pressure chosen together. Bioink delivery of 1 µL/s requires 72.7 kPa for a 200 µm diameter nozzle, producing a D_t_ of 5.5%, but it only requires 26.6 kPa for a 330 µm diameter nozzle, producing a D_t_ of 4.6% (Fig. 7). At a fixed flow rate of 770 µL/s and nozzle outlet diameters of 0.50 to 1.00 mm, wider than any modeled here, Lombardi *et al*. report the opposite ordering, with damage increasing with outlet diameter [14].

### Extensional dominance and its design implications

Because the entrance term carries 97.1% of the predicted damage at the reference condition, the abrupt step from the cartridge outlet into the nozzle is a geometrical feature to be potentially redesigned. The converging streamlines carry an extensional strain rate that peaks at 5.44 × 10^3^ s^−1^ over an axial extent of about 130 µm (Fig. 8). A tapered or hyperbolically streamlined transition is expected to reduce p_en_ and D_et_ without any change in nozzle diameter or applied pressure [7,14]. Ning *et al*. likewise recommend tapered needles, which reach a given flow rate at a lower pressure and so lower both the shear and the extensional stress [9]. Lombardi *et al*. related the balance of shear and extensional stress to measured viability when they varied the contraction angle experimentally. For red blood cells, in a multilaboratory study of the benchmark nozzle of the U.S. Food and Drug Administration at a matched flow rate of 6 L/min, a sudden-contraction inlet produced a modified index of hemolysis of 1.239 ± 0.667, compared with 0.021 ± 0.128 for a gradual cone [33].

A streamlined transition acts on the viscous converging-flow loss [34–36], not on the inertial part of the entrance pressure, which is the kinetic energy of the jet. That part is 6.6% of the residual at the reference condition (200 kPa, 200 µm diameter nozzle), but 59.1% for a 330 µm diameter nozzle at 200 kPa (Additional file 1: Table S1), which limits what a taper can remove at the widest nozzles.

### Comparison with prior computational approaches

Among the mechanistic bioprinting simulations compared in Table 4, the present model is the only one that combines a damage law calibrated for the printed cell line with a surrogate of the pressure–diameter plane. For instance, Reina-Romo *et al*. coupled computational fluid dynamics (CFD) with a GP to screen nozzle designs against generic shear-stress thresholds [7]. McCauley *et al*. identify cell deformability and viscoelastic relaxation as factors that a power-law model does not capture [10]. Chirianni *et al*. built operative nomograms of flow rate and viability over printing pressure and nozzle diameter, combining CFD of the same contraction-plus-capillary topology with a reduced-order model and a generalized logistic damage law. They resolved the pressure drop across the contraction directly, whereas the present entrance pressure is the residual of the applied pressure and the developed nozzle drop [8]. Friedrich *et al*. found, in simulations of embedded bioprinting, that a conical nozzle reduces damage in cells sensitive to the magnitude of stress but can increase it in cells sensitive to its duration [16]. Müller *et al*. coupled lattice Boltzmann flow with a finite element cell model and showed that, for cells near the nozzle center, the extensional stress at the nozzle exit can be significant [17].

**Table 4.** Comparison of the present pipeline with five prior mechanistic bioprinting simulations.

| Study | Ref. | Flow solver | Particle tracking | Damage model | Principal finding | Cell type |
| --- | --- | --- | --- | --- | --- | --- |
| Present work |  | Axisymmetric power-law Navier–Stokes, FEM (DOLFINx 0.10) | Lagrangian, 168 tracers, RK45 | Ning *et al.* shear–extensional cascade, L8-calibrated | +101.0% total damage, Ø200 to Ø330 µm at fixed 200 kPa; order reverses at matched flow rate. | L8 myoblasts in alginate |
| Reina-Romo *et al.* 2021 | [7] | CFD (OpenFOAM) | None; Eulerian streamlines | Generic shear-stress threshold | Middle and exit *radii* and nozzle length set peak shear; geometry and shear-thinning act together. | Generic shear-thinning hydrogel |
| McCauley *et al.* 2025 | [10] | Analytical and CFD | None | Critical-strain model | Viability set by coupled process, rheology and cell parameters. | Generic cell types |
| Friedrich *et al.* 2022 | [16] | CFD, finite volume (OpenFOAM) | Eulerian pathlines at fixed *radii* | Ning *et al.*, cumulative survival | Magnitude- and duration-sensitive regimes; one-dimensional radial profiles. | Synthetic parameters |
| Müller *et al.* 2023 | [17] | Lattice Boltzmann with FEM cell | Immersed boundary, cell-resolved | Hyperelastic (neo-Hookean) | Extensional stress at the nozzle exit affects centerline cells. | Generic |
| Chirianni *et al.* 2024 | [8] | CFD | Two representative inlet *radii* | Generalized logistic law, not derived from the Ning law | Extensional stress exceeds shear in the contraction; damage nomograms are presented. | Generic |

The present model adds a decomposition of the diameter response for L8 myoblasts into extensional, shear, friction and inertial parts on one 32-case grid, and an operating window tested at eleven conditions solved after the surrogate had fixed them.

### Computational cost of the surrogate

Once trained, the surrogate evaluates a condition in 0.08 ms and searches the full operating window, 3751 pressures at each of the four diameters, in 0.14 s (Additional file 1: Section S4). Solving the same 15 004 conditions with the finite element model would take about 22 000 CPU-hours, compared with the 47 CPU-hours of the 32-case training campaign. Direct interpolation of the solved cases recovers the damage at the operating points of the four solved diameters to within 0.02 percentage points. What the surrogate adds is the diameters that were not solved, checked at 150 and 290 µm, and a map of its own uncertainty (Fig. 6D). Interpolating a deterministic simulation campaign with a GP is the standard treatment of such computer experiments [37,38], and the posterior standard deviation could direct adaptive sampling toward conditions where high predicted D_t_ meets high uncertainty.

### Comparison with measured flow and damage

The model follows the rise of the damage that Ning *et al*. measured in L8 skeletal muscle cells immediately after printing through the 200 µm diameter nozzle, and underestimates it by 2.1 to 6.6 percentage points. At 50, 100, 200 and 400 kPa, it predicts 4.89, 6.28, 8.96 and 20.53%, compared with 6.99, 11.70, 15.53 and 23.08% measured. These are the conditions on which Ning *et al*. fitted a_2_ and b_2_, to the measured damage minus their own shear estimate of about 5% [9].

Following Eq. 21, the same shear law integrated over the developed nozzle profile gives 0.16 to 0.95%. Their estimate exceeds this integral by 4.9 to 5.1 percentage points at every pressure, and by 5.0 to 5.2 in their RSC96 cells, although the integral falls sixfold between 50 and 400 kPa. Evaluated on a discrete radial grid, the area average counts the wall node, where the velocity is zero and the exposure time unbounded, as fully damaged, which adds a term that does not depend on pressure. A grid of 20 intervals reproduces the four shear-damage estimates of Ning *et al*. for L8 cells in this way with a root-mean-square difference of 0.12 percentage points (Additional file 1: Section S3). The model therefore lies below the measurements by about 5 percentage points, the amount assigned to shear before the extensional branch was fitted, which neither branch of the law reproduces once the shear integral is evaluated exactly. The remainder, −2.8 to +1.5 percentage points, spans the same range as the residuals of that fit.

The flow-rate over-prediction has the same sign at all five pressure values (Fig. 4A). The wall shear stresses Ning *et al*. report imply, through Eq. 16, flow rates within 1.4% of the present ones for the 200 µm diameter nozzle (Table 1, Fig. 4B), so their simulation carries the same offset. The shared offset points to the inputs common to both models, the rheology and the geometry.

Because the entrance pressure is a small residual of two large pressures, the extensional damage is sensitive to the flow rate: at 200 kPa, the measured flow rate raises it by 3.1 percentage points, from 8.69 to 11.8%. Because every operating point lies on the flow limit, a true flow rate below the prediction would also raise the operating pressures. The residual entrance pressure exceeds the entrance pressures of Ning *et al*. by 0.6 to 6.7% across the four diameters at 200 kPa, differences of the order documented between entrance-pressure extraction methods in capillary rheometry [39].

### Limitations and future work

The power-law model omits yield stress, viscoelasticity and thixotropy, which Herschel– Bulkley, viscoelastic and thixotropic descriptions would capture for many practical bioinks [10,30]. The extensional term is also uniform within each case; integrating extensional strain along individual trajectories through the contraction would give a particle-resolved D_et_.

As in the model of Ning *et al*., which assigns shear damage to the nozzle, shear exposure upstream of the nozzle is excluded. At the reference condition, the residence times are 8.11 s in the cartridge and 0.66 s in the truncated-cone transition, compared with 0.018 s in the nozzle. Applied to these two regions, the shear law would add 1.13 percentage points to D_st_, about four times the 0.29 percentage points accumulated in the nozzle. It would do so at a wall shear stress of 40.1 Pa, below the 100 Pa lower bound of its calibration [9]. Including this term would raise the total damage and reduce the increase in damage from the 200 to the 330 µm diameter nozzle at 200 kPa without changing its direction.

In 13 of the 32 cases, the effective extensional stress lies outside the range on which the extensional branch was calibrated (Table 3; Additional file 1: Table S1). The 20% damage limit also lies just above its calibrated range, since Eq. 23 gives D_et_ = 19.5% at 9251 Pa. Recalibrating the law over a wider stress range would remove this restriction, and the coefficients, specific to L8 myoblasts, must be recalibrated for other cell lines [9]. Printing L8 cells through two nozzle diameters, once at fixed pressure and once at matched flow rate, would test the constraint-dependent ordering directly in this cell line.

## Conclusions

For L8 myoblasts extruded in a 1.5% alginate suspension, a finite element–Gaussian process surrogate predicts flow rate and mechanical damage for any nozzle diameter and pressure within the simulated range in under a millisecond. One finite element case takes about 1.5 hours, so printing parameters that meet a throughput target and a damage limit can be selected in real time, without new simulations. The simulations also show that the predicted damage comes mainly from the entrance to the nozzle. At fixed pressures of 75 kPa or more, a wider nozzle is more damaging, while at matched flow rate the widest nozzle is the least damaging, although the differences are small. Nozzle diameter and pressure should therefore be chosen together for the flow rate the print requires, and the nozzle geometry may be the feature to redesign to protect cell viability.

## Supplementary Information

Additional file 1 (.docx): Supplementary methods, Figures S1–S3 and Tables S1–S5. Viscosity regularization and weak form; damage, calibration ranges and inertial share for all 32 cases; surrogate implementation and held-out validation; mesh independence and Bagley check; interval calibration and sensitivity of the surrogate; FEM checks of the operating window.

Additional file 2 (.mp4): Velocity field through the cartridge-to-nozzle contraction at ΔP = 200 kPa, 200 µm diameter nozzle, with 900 visualization particles.

## List of abbreviations

CFD: computational fluid dynamics
FEM: finite element modeling
GP: Gaussian process
ML: machine learning
RK45: explicit Runge–Kutta method of order 5(4)
D_t_: total predicted mechanical damage
D_st_: area-averaged shear damage in the nozzle
D_et_: extensional damage acquired at the nozzle entrance
p_en_: entrance pressure
Re: Reynolds number

## Declarations

### Ethics approval and consent to participate

Not applicable.

### Consent for publication

Not applicable.

### Availability of data and materials

The simulation data and code supporting this study are available in the GitHub repository https://github.com/tavangariz/l8-myoblast-bioprinting-fem, archived on Zenodo as release v1.0.0 (https://doi.org/10.5281/zenodo.23469527).

### Authors’ contributions

ZT: Conceptualization, methodology, software, formal analysis, investigation, writing – original draft, visualization.

AM: Methodology, software, validation, writing – review and editing. AP: Supervision, writing – review and editing.

GGG: Conceptualization, supervision, writing – review and editing, funding acquisition.

All authors read and approved the final manuscript.

## Acknowledgements

The authors gratefully acknowledge ASI for funding of this study. The authors also thank the ReCaS-Bari data center for access to and support with the high-throughput computing infrastructure on which the finite element campaign was run.

