## Additional file 1 for "Fast computational screening of extrusion parameters for bioprinting of skeletal myoblasts in alginate hydrogel"

#### S1 Numerical regularization

The singular power-law viscosity and the axisymmetric terms were regularized for nonlinear solution robustness. The radial denominator was replaced by

$$r_\varepsilon = r + \varepsilon_r, \quad \varepsilon_r = 10^{-12}\ \mathrm{m} \tag{S1}$$

and the shear-rate invariant used in the finite element implementation was

$$\dot{\gamma}_\varepsilon = \left[2\left(\boldsymbol{D}_{2\mathrm{D}} : \boldsymbol{D}_{2\mathrm{D}} + \left(\frac{u_r}{r_\varepsilon}\right)^2\right) + \varepsilon_{\dot{\gamma}}^2\right]^{1/2}, \quad \varepsilon_{\dot{\gamma}}^2 = 10^{-8}\ \mathrm{s}^{-2} \tag{S2}$$

where

$$\boldsymbol{D}_{2\mathrm{D}} = \mathrm{sym}\left(\nabla_{r,z}\boldsymbol{u}\right) \tag{S3}$$

contains the in-plane $(r, z)$ components. The power-law viscosity was smoothly bounded according to

$$\eta_\varepsilon = \mathrm{smin}_{\delta_\eta}\left[\mathrm{smax}_{\delta_\eta}\left(K\dot{\gamma}_\varepsilon^{\,n-1}, \mu_{\mathrm{min}}\right), \mu_{\mathrm{max}}\right] \tag{S4}$$

where

$$\mathrm{smax}_\delta(a, b) = \frac{1}{2}\left[a + b + \sqrt{(a - b)^2 + \delta^2}\right] \tag{S5}$$

$$\mathrm{smin}_\delta(a, b) = \frac{1}{2}\left[a + b - \sqrt{(a - b)^2 + \delta^2}\right] \tag{S6}$$

The final viscosity bounds were

$$\mu_{\mathrm{min}} = 0.003\ \mathrm{Pa \cdot s}, \quad \mu_{\mathrm{max}} = 500\ \mathrm{Pa \cdot s}, \quad \delta_\eta = 10^{-3}\ \mathrm{Pa \cdot s} \tag{S7}$$

The lower bound was reduced through the continuation sequence 10, 1, 0.1, 0.01 and 0.003 Pa·s, with the solution at each level used to initialize the next nonlinear solve. At the first viscosity level, the applied pressure was increased in five equal steps from $\Delta P/5$ to $\Delta P$.

The meshes were generated with Gmsh and refined anisotropically toward the nozzle: the nozzle radius carries twelve elements (mesh M12, elements of 4.2–13.8 µm, scaling as R/12), against roughly 200 µm *per* element in the cartridge, and the mesh carries 47 952, 22 369, 17 602 and 13 584 nodes within 100, 200, 250 and 330 µm diameter nozzles respectively. The nonlinear system was solved by a Newton iteration implemented directly (PETSc MUMPS LU, 8-step backtracking line search, up to 40 iterations, rtol = $10^{-6}$, atol = $10^{-8}$). Jobs were executed on the ReCaS high-throughput computing cluster (HTCondor, single-process Apptainer mode) at 1.5 hours of compute *per* case on average (0.9 to 2.3 hours across the 32 cases).

Because wall shear stress is formed from the solved flow rate through the Rabinowitsch relation, the regularization enters the reported quantities only through the flow field. At the analytic wall shear rate, the $10^{-8}$ $s^{-2}$ floor changes the shear rate by less than $10^{-10}$ relative, and the bounded viscosity differs from $K\dot{\gamma}^{n-1}$ by 0.03% for a 200 µm diameter nozzle at 200 kPa, and by 1.4% for a 330 µm diameter nozzle at 400 kPa, where the wall viscosity approaches its lower bound.

#### S2 Finite element weak form

The equations were discretized on the meridional ($r$, $z$) domain using Taylor–Hood elements, with continuous quadratic interpolation for velocity and continuous linear interpolation for pressure. With $v$ and $q$ the velocity and pressure test functions, the implemented residual at the final continuation level was

$$0 = \int_{\Omega} 2\pi r \left\{\rho\left[\left(\boldsymbol{u} \cdot \nabla_{r,z}\right)\boldsymbol{u}\right] \cdot \boldsymbol{v} + 2\eta_{\varepsilon}\left[\boldsymbol{D}_{2\mathrm{D}}(\boldsymbol{u}) : \boldsymbol{D}_{2\mathrm{D}}(\boldsymbol{v}) + \frac{u_r v_r}{r_{\varepsilon}^2}\right] - p\left(\nabla_{r,z} \cdot \boldsymbol{v} + \frac{v_r}{r_{\varepsilon}}\right) - q\left(\nabla_{r,z} \cdot \boldsymbol{u} + \frac{u_r}{r_{\varepsilon}}\right)\right\} \mathrm{d}A + \int_{\Gamma_{\mathrm{in}}} 2\pi r\, \Delta P\, \boldsymbol{n} \cdot \boldsymbol{v}\, \mathrm{d}s \qquad \text{(S8)}$$

The factor $2\pi r$ converts the meridional-area and boundary integrals to their three-dimensional axisymmetric counterparts. The nonlinear residual and its consistent Jacobian were assembled automatically and solved by Newton iteration. The convective term of Eq. S8 carries a density continuation factor. Inertia was introduced at the final viscosity level with ρ-fractions 0.1, 0.25, 0.5, 0.75 and 1.0, and a case was kept only if the Newton solve at ρ-fraction 1 converged.

#### S3 Damage, calibration ranges and inertial share for every case

The shear damage of the cells collected at the outlet weights the local damage of Eq. 20 of the main text by the axial velocity of Eq. 19, where Eq. 21 weights it by area:

$$D_{\mathrm{st,Q}} = \frac{2\pi}{Q}\int_0^R r\, v_z(r)\, D_{\mathrm{s}}(r)\, \mathrm{d}r \qquad \text{(S9)}$$

where $Q$ is the volumetric flow rate and $R$ the nozzle radius. Table S1 lists both averages for every case; the area average is the convention of Ning *et al.* (main text, reference 9) and the one used in the main text. The two averages are at most 1.46 percentage points apart, for the 100 μm diameter nozzle at 25 kPa.

Across the grid, $\tau_e$ ranges from 101 to 59 727 Pa, with the widest nozzle at 400 kPa exceeding the 9251 Pa upper limit of the extensional branch by a factor of 6.5. The shear branch was fitted on exposure times of 10 to 120 s, which sit above most of this grid: nozzle residence spans 11.5 to 135 ms across the tracked radius at the reference condition. Over the whole grid, the mean residence time, $L_{\mathrm{tip}}/\bar{u}$, ranges from 0.90 ms for the 330 μm diameter nozzle at 400 kPa to

93.6 s for the 100 μm diameter nozzle at 25 kPa, and only this nozzle, at 25 and 50 kPa (93.6 and 12.9 s), has a residence time inside the fitted window. The largest $D_{st}$ in the grid, 4.34 percentage points for the 100 μm diameter nozzle at 25 kPa, is an extrapolation below the fitted stresses: its residence time lies inside the fitted window, but its wall stress, $\tau_w = 62$ Pa, lies below the fitted range. The shear exponents ($b_1 = 0.61$, $c_1 = 0.52$) were checked against two benchmark values reproduced from Ning *et al.* Part of the nozzle cross-section lies inside the calibrated shear range in 22 of the 32 cases (Table S1).

***Table S1*** *Damage, shear averages, calibration ranges and inertial share for all 32 combinations of nozzle diameter and applied pressure*

| Ø (μm) | $\Delta P$ (kPa) | $D_t$ (%) | $D_{st}$ area (%) | $D_{st,Q}$ flow (%) | In shear range (%) | $\tau_e$ (Pa) | $p_{en}$ (kPa) | Inertial share (%) |
|---|---|---|---|---|---|---|---|---|
| 100 | 25 | 6.05 | 4.336 | 2.873 | 0 | 101† | 0.20 | 0.0 |
| 100 | 50 | 4.95 | 2.425 | 1.579 | 15.8 | 198† | 0.39 | 0.0 |
| 100 | 75 | 4.92 | 1.718 | 1.111 | 3.1 | 303† | 0.60 | 0.0 |
| 100 | 100 | 5.14 | 1.344 | 0.865 | 0.6 | 413† | 0.82 | 0.0 |
| 100 | 150 | 5.75 | 0.949 | 0.607 | <0.1 | 634 | 1.25 | 0.0 |
| 100 | 200 | 6.36 | 0.740 | 0.473 | <0.1 | 847 | 1.67 | 0.0 |
| 100 | 300 | 7.55 | 0.521 | 0.332 | <0.1 | 1284 | 2.54 | 0.3 |
| 100 | 400 | 8.73 | 0.406 | 0.258 | <0.1 | 1766 | 3.49 | 1.2 |
| 200 | 25 | 4.35 | 1.722 | 1.113 | 8.8 | 210† | 0.42 | 0.0 |
| 200 | 50 | 4.89 | 0.951 | 0.609 | 0.1 | 439† | 0.87 | 0.0 |
| 200 | 75 | 5.59 | 0.670 | 0.428 | <0.1 | 659 | 1.30 | 0.0 |
| 200 | 100 | 6.28 | 0.522 | 0.333 | <0.1 | 882 | 1.74 | 0.3 |
| 200 | 150 | 7.59 | 0.368 | 0.233 | <0.1 | 1347 | 2.66 | 1.7 |
| 200 | 200 | 8.96 | 0.286 | 0.182 | <0.1 | 1902 | 3.76 | 6.6 |
| 200 | 300 | 13.09 | 0.203 | 0.128 | 0 | 4069 | 8.04 | 31.7 |
| 200 | 400 | 20.53 | 0.161 | 0.102 | 0 | 10 125† | 20.00 | 58.9 |
| 250 | 25 | 4.35 | 1.276 | 0.821 | 1.5 | 279† | 0.55 | 0.0 |
| 250 | 50 | 5.21 | 0.703 | 0.448 | <0.1 | 559 | 1.11 | 0.0 |
| 250 | 75 | 6.11 | 0.495 | 0.315 | <0.1 | 841 | 1.66 | 0.2 |
| 250 | 100 | 6.97 | 0.385 | 0.245 | <0.1 | 1133 | 2.24 | 1.0 |
| 250 | 150 | 8.75 | 0.271 | 0.172 | <0.1 | 1820 | 3.60 | 7.1 |
| 250 | 200 | 11.16 | 0.212 | 0.134 | 0 | 2969 | 5.87 | 23.1 |
| 250 | 300 | 20.05 | 0.154 | 0.097 | 0 | 9648† | 19.06 | 61.5 |
| 250 | 400 | 32.02 | 0.127 | 0.081 | 0 | 26 159† | 51.67 | 76.1 |
| 330 | 25 | 4.53 | 0.879 | 0.562 | 0.2 | 380† | 0.75 | 0.0 |
| 330 | 50 | 5.81 | 0.482 | 0.307 | <0.1 | 762 | 1.50 | 0.3 |
| 330 | 75 | 7.01 | 0.339 | 0.215 | <0.1 | 1161 | 2.29 | 1.7 |
| 330 | 100 | 8.25 | 0.265 | 0.168 | <0.1 | 1628 | 3.21 | 6.3 |
| 330 | 150 | 11.85 | 0.188 | 0.119 | 0 | 3355 | 6.63 | 31.1 |
| 330 | 200 | 18.00 | 0.151 | 0.095 | 0 | 7746 | 15.30 | 59.1 |
| 330 | 300 | 33.06 | 0.118 | 0.075 | 0 | 28 126† | 55.56 | 79.5 |
| 330 | 400 | 45.49 | 0.104 | 0.066 | 0 | 59 727† | 117.98 | 78.5 |

*$D_t$: total damage. $D_{st}$: shear damage averaged over the nozzle cross-section by area, see Eq. 21 of the main text and the convention of Ning et al. $D_{st,Q}$: the same average weighted by the local axial velocity (flow-weighted), see Eq. S9. In shear range: the flow-weighted fraction of the nozzle cross-section in which the local shear stress and exposure time lie jointly inside the calibrated range of the shear branch, 100 to 1000 Pa and 10 to 120 s. $\tau_e$: effective extensional stress (main text, Methods). † marks the 13 values outside the calibrated 440 to 9251 Pa of the extensional branch (8 below, 5 above). $p_{en}$: entrance pressure. Inertial share: the fraction of $p_{en}$ removed when the same case is solved without the convective term (main text, Results). M12 mesh applies throughout.*

The shear damage that Ning *et al.* plot for their 200 µm diameter nozzle is 5.82, 5.55, 5.36 and 5.16% for L8 cells at 50, 100, 200 and 400 kPa, and 5.68, 5.43, 5.21 and 5.19% for RSC96 cells. Their Eq. 9, the area average of the local shear damage over the nozzle cross-section, evaluated with their shear laws, rheology and Table 2 (Extensional and shear stresses under varying bioprinting conditions) entrance pressures, gives 0.951, 0.522, 0.286 and 0.160% for L8 cells, equal to the integral of Eq. 21 of the main text evaluated with their inputs, and 0.643, 0.245, 0.092 and 0.036% for RSC96 cells. The difference, 4.9 to 5.1 percentage points for L8 cells and 5.0 to 5.2 for RSC96 cells, barely changes with pressure, while the integral falls six- and eighteen-fold. On a discrete radial grid the wall node, where the velocity is zero and the exposure time unbounded, carries $D_s$ = 100%, and with the trapezoid rule on $N$ intervals that node alone adds 100/$N$ percentage points. With 20 intervals, the rule reproduces the four values for L8 cells to within 0.12 percentage points (root mean square), and with 19 the values for RSC96 cells to within 0.07 percentage points.

#### S4 Surrogate implementation and validation

The implementation is scikit-learn's GaussianProcessRegressor, with the constant variance bounded between $10^{-3}$ and $10^{3}$ and the length scales between $10^{-2}$ and $10^{2}$ in standardized units, normalized targets and random seed 0. The Matérn 5/2 kernel assumes a twice-differentiable response; that assumption is tested against held-out data rather than inferred from the smoothness of the fitted surface. The ten restarts of the optimizer are repeated attempts to reach the best optimum, not validation folds. The fitted kernels are $2.27^2$ × Matérn(3.07, 3.44) for $D_t$, $21.3^2$ × Matérn(6.13, 21.5) for $\log_{10} Q$ and $18.6^2$ × Matérn(5.16, 24.1) for $\log_{10} \tau_w$, with length scales for pressure and diameter in that order. None is at a bound of its search range, and the final fits raised no optimizer convergence warning, whereas the refits inside the held-out folds raised 18, 6 and 2 such warnings with single cases, diameters and pressures held out.

Leaving out one case at a time, the mean absolute error in flow rate was 0.027 in $\log_{10} Q$, the root-mean-square error 0.054 and the largest error 0.251. When computed directly from the held-out predictions, the mean absolute percentage error in $Q$ was 6.8%. In wall shear stress, the mean absolute error was 0.006, the root-mean-square error 0.013 and the largest 0.059, all in $\log_{10} \tau_w$.

Fitting the three GPs took 1.0 s, the median of 5 repeats, and a single query 0.08 ms, the median of 2000 repeated queries. The full operating-window search, 3751 pressures at each of the four solved diameters for two outputs in one batch of 15 004 points, takes 0.14 s, the median of 50 repeats. Both were timed on one Intel Core Ultra 9 288V laptop processor with Python 3.13.9, NumPy 2.2.5, SciPy 1.15.3 and scikit-learn 1.7.2.

Fig. S1 and Table S2 give the error for every held-out nozzle diameter and applied pressure. With a whole diameter held out, the largest single error in $D_t$ is 3.47 percentage points. Across the four held-out diameters, the root-mean-square error in total damage is 1.21 percentage points. In flow rate, the logarithmic error is 0.169 on average, 0.207 root-mean-square and 0.403 at most, an error of 47% on average and 153% at most in the held-out predictions. In wall shear stress, it is 0.028, 0.035 and 0.076. Across the eight held-out pressures, the root-mean-square error in total damage is 1.13 percentage points and the largest 4.05. In flow rate, the logarithmic error is 0.124

on average, 0.199 root-mean-square and 0.614 at most, 24% and 76% in the held-out predictions. In wall shear stress, it is 0.049, 0.081 and 0.240.

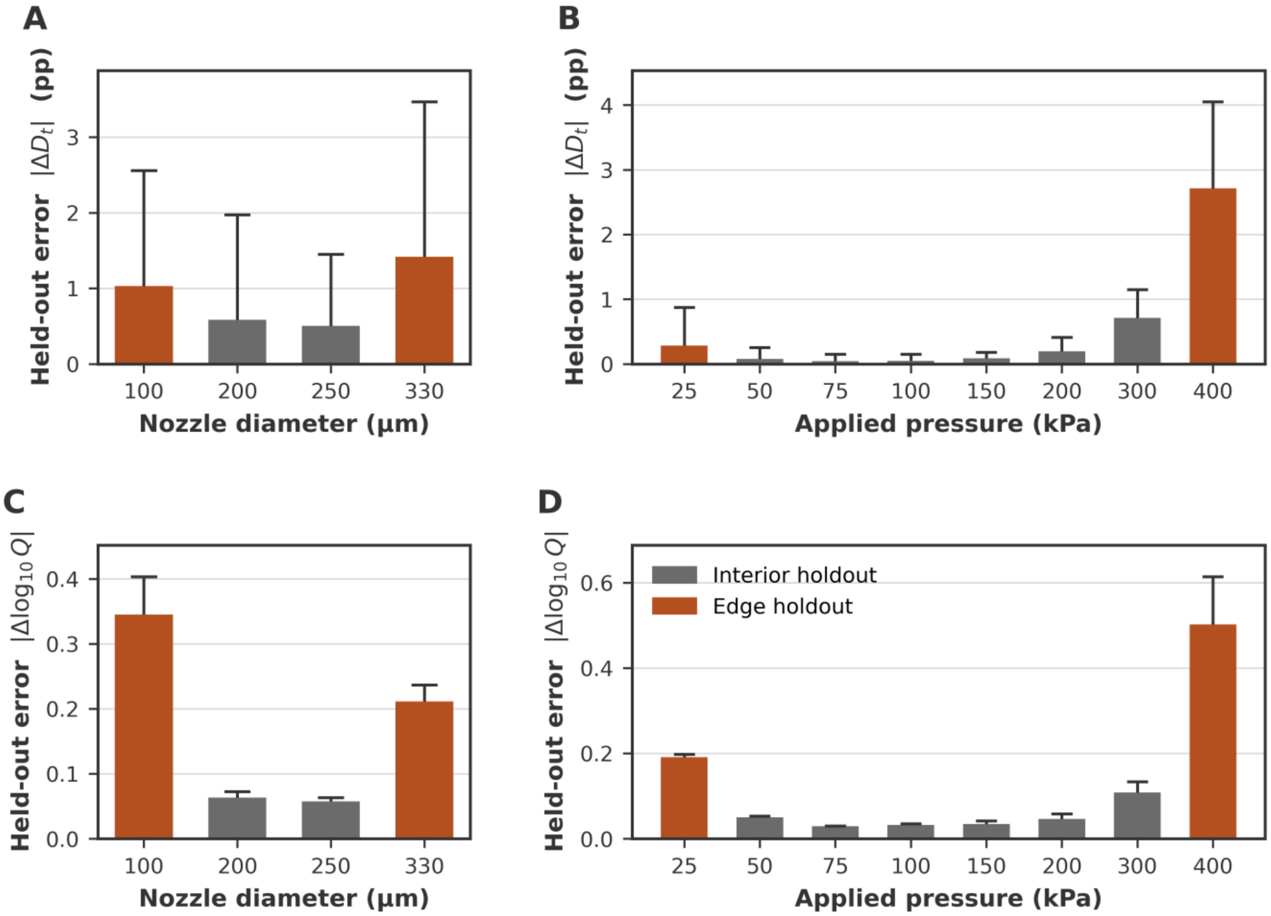


***Fig. S1*** *Surrogate error with a whole nozzle diameter or a whole applied pressure held out. (A) $D_t$ error against the held-out nozzle diameter. (B) $D_t$ error against the held-out applied pressure. (C, D) The same for $\log_{10} Q$. Bars show the mean absolute error over the cases held out together, while caps mark the largest error. Gray bars are interior holdouts, which test interpolation across a missing level; orange bars are edge holdouts, which test extrapolation beyond the remaining levels. Every prediction comes from a model refitted, standardization included, without the held-out level.*

Leaving out one case at a time, the exact binomial interval on the coverage of the nominal 95% interval is 67 to 95%, and with 32 dependent cross-validation predictions the degree of undercoverage is itself uncertain. With a whole diameter held out, the intervals on $D_t$ widened, from a mean width of 1.7 to 14.2 percentage points, and contained all 32 values, so full coverage there reflects the wide intervals; with a pressure held out they averaged 3.3 percentage points and contained 29 of 32. The intervals on $\log_{10} Q$ contained all 32 values with single cases held out, at a mean width of 0.136, but widened only to 0.157 with a diameter held out and then contained none, at the interior diameters as well as the edges; with a pressure held out they widened to 1.034 and contained 20 of 32.

***Table S2*** *Surrogate error for every held-out nozzle diameter and applied pressure*

| Held out | Level | Type | $D_t$ MAE (pp) | $D_t$ max (pp) | $\log_{10} Q$ MAE | $\log_{10} Q$ max | $D_t$ in 95% interval | $\log_{10} Q$ in 95% interval |
|---|---|---|---|---|---|---|---|---|
| Diameter | 100 µm | edge | 1.030 | 2.557 | 0.345 | 0.403 | 8/8 | 0/8 |
| Diameter | 200 µm | interior | 0.582 | 1.975 | 0.063 | 0.073 | 8/8 | 0/8 |
| Diameter | 250 µm | interior | 0.504 | 1.450 | 0.057 | 0.063 | 8/8 | 0/8 |
| Diameter | 330 µm | edge | 1.415 | 3.465 | 0.211 | 0.237 | 8/8 | 0/8 |
| Pressure | 25 kPa | edge | 0.284 | 0.875 | 0.191 | 0.198 | 3/4 | 0/4 |
| Pressure | 50 kPa | interior | 0.079 | 0.256 | 0.050 | 0.053 | 3/4 | 0/4 |
| Pressure | 75 kPa | interior | 0.045 | 0.153 | 0.029 | 0.030 | 3/4 | 0/4 |
| Pressure | 100 kPa | interior | 0.046 | 0.150 | 0.032 | 0.035 | 4/4 | 4/4 |
| Pressure | 150 kPa | interior | 0.089 | 0.179 | 0.035 | 0.042 | 4/4 | 4/4 |
| Pressure | 200 kPa | interior | 0.196 | 0.414 | 0.047 | 0.059 | 4/4 | 4/4 |
| Pressure | 300 kPa | interior | 0.712 | 1.150 | 0.108 | 0.134 | 4/4 | 4/4 |
| Pressure | 400 kPa | edge | 2.712 | 4.049 | 0.502 | 0.614 | 4/4 | 4/4 |

MAE, mean absolute error over the cases held out together; max, the largest of them; pp, percentage points. Interior levels test interpolation across a missing level and edge levels extrapolation beyond the remaining ones. The last two columns count held-out values inside the nominal 95% interval of the GP posterior. Fold-level predictions: fold_predictions.csv in the results/surrogate folder of the code repository.

Table S3 compares the surrogate with linear, quadratic and cubic polynomial surfaces refitted on the same held-out splits.

***Table S3*** *Surrogate and polynomial baselines on identical held-out splits*

| Model | Case held out (pp) | Diameter held out (pp) | Pressure held out (pp) |
|---|---|---|---|
| Gaussian process, Matérn 5/2 | 0.338 | 0.883 | 0.520 |
| Linear surface | 4.242 | 4.914 | 4.229 |
| Quadratic surface | 1.273 | 2.070 | 1.276 |
| Cubic surface | 0.931 | 2.986 | 0.840 |

Values are the mean absolute error in $D_t$, in percentage points, with every model refitted on the same training rows in every fold. Polynomial surfaces are least-squares fits in applied pressure and nozzle diameter.

### S5 Model checks

Fig. S2 collects three checks on the model: mesh independence between the M12 and M16 meshes, the entrance share of the predicted damage across the grid, and the entrance pressure against a Bagley extrapolation to zero nozzle length.

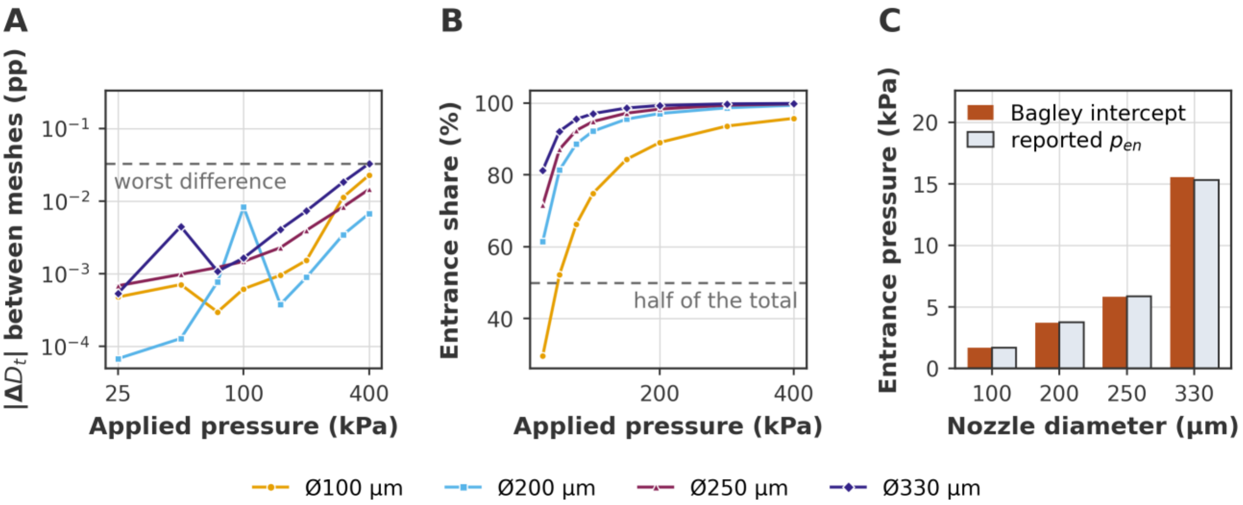


***Fig. S2*** *Three supporting checks on the model. (A) Mesh independence: the absolute change in predicted total damage between the M12 and M16 meshes; the dashed line marks the largest difference anywhere in the grid, 0.033 percentage points. (B) The entrance share of the predicted damage,* $D_{et}/D_t$*, against applied pressure for the four nozzles. (C) The entrance pressure the damage law consumes against an independent Bagley extrapolation to zero nozzle length: the two agree to better than 2% at every diameter.*

#### S6 Interval calibration and sensitivity of the surrogate

Of the 32 held-out predictions of $D_t$, leaving out one case at a time, 27 fall inside their own nominal 95% interval, having a half-width of 1.96 posterior standard deviations. Fig. S3 plots each error against that half-width.

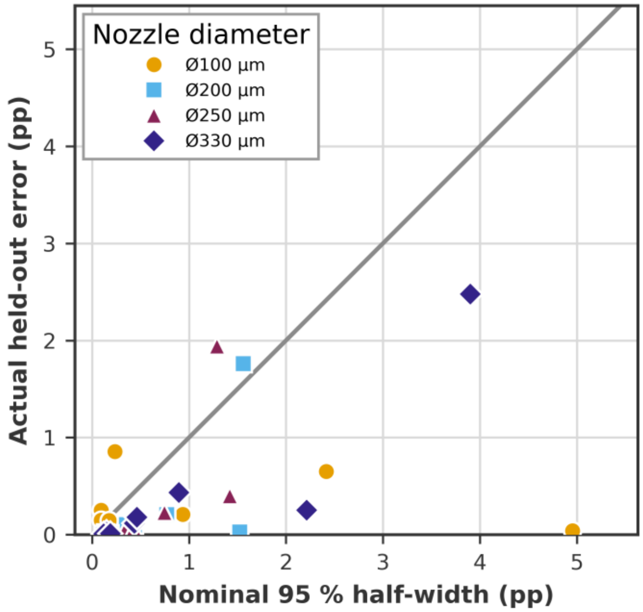


***Fig. S3*** *Held-out error against the nominal 95% interval of the surrogate, leaving out one case at a time. Each point is one of the 32 predictions of $D_t$, plotted at its nominal 95% half-width against its actual error; a point below the line fell inside its own interval. The interval is the conditional GP posterior, not a calibrated error bar.*

Table S4 repeats the three held-out tests with a rougher Matérn 3/2 kernel and with the regularization α raised to $10^{-4}$ and $10^{-2}$.

***Table S4*** *Sensitivity of the surrogate to kernel smoothness and the regularization α*

| Kernel | α | $D_t$ MAE, case (pp) | $D_t$ MAE, diameter (pp) | $D_t$ MAE, pressure (pp) | $D_t$ coverage, case (%) | $\log_{10} Q$ coverage, diameter (%) | Operating pressure at Ø100 / 200 / 250 / 330 µm (kPa) |
|---|---|---|---|---|---|---|---|
| Matérn 5/2 | $10^{-6}$ | 0.338 | 0.883 | 0.520 | 84.4 | 0.0 | 298.5 / 72.7 / 46.5 / 26.6 |
| Matérn 5/2 | $10^{-4}$ | 0.343 | 1.034 | 0.457 | 90.6 | 15.6 | 298.4 / 72.4 / 46.7 / 26.6 |
| Matérn 5/2 | $10^{-2}$ | 0.668 | 1.640 | 0.598 | 96.9 | 71.9 | 294.5 / 77.3 / 49.0 / 25.0 |
| Matérn 3/2 | $10^{-6}$ | 0.348 | 2.359 | 0.357 | 93.8 | 75.0 | 298.6 / 72.6 / 46.6 / 26.8 |
| Matérn 3/2 | $10^{-4}$ | 0.358 | 2.406 | 0.363 | 93.8 | 75.0 | 298.4 / 72.5 / 46.7 / 26.7 |
| Matérn 3/2 | $10^{-2}$ | 0.528 | 2.272 | 0.625 | 100.0 | 75.0 | 291.1 / 76.2 / 49.1 / 25.0 |

*The held-out scheme is named by case, diameter or pressure, and coverage is the percentage of held-out values inside the nominal 95% interval. The operating pressure is the lowest-$D_t$ pressure in 25 to 400 kPa satisfying $D_t < 20\%$ and $Q \geq 1$ µL/s. The first row is the configuration used throughout the paper; the others were computed after it, and none was selected from them.*

### S7 FEM checks of the operating window

Eleven conditions were fixed from the surrogate predictions before any of them was solved (main text, Methods) and solved with the same model and settings as the grid. Table S5 compares each with the surrogate, and they are depicted as diamonds in Fig. 7A (main text).

***Table S5*** *Surrogate predictions compared with eleven FEM checks fixed before solving*

| Ø (µm) | $\Delta P$ (kPa) | Condition | $D_t$, FEM (%) | $D_t$, surrogate (%) | Error (pp) | $Q$, FEM (µL/s) | $Q$ error (%) | In window, FEM / surrogate |
|---|---|---|---|---|---|---|---|---|
| 100 | 298.5 | operating point | 7.53 | 7.54 | 0.00 | 1.001 | −0.1 | yes / yes |
| 150 | 133.2 | operating point | 6.29 | 6.13 | −0.16 | 1.060 | −5.5 | yes / yes |
| 150 | 300.0 | interior | 9.50 | 9.09 | −0.40 | 10.74 | −5.7 | yes / yes |
| 200 | 72.7 | operating point | 5.53 | 5.52 | 0.00 | 0.9998 | +0.3 | no / yes |
| 200 | 393.4 | $D_t$ = 20% | 19.95 | 20.01 | +0.06 | 113.7 | +2.1 | yes / no |
| 250 | 46.5 | operating point | 5.08 | 5.08 | 0.00 | 1.016 | −1.1 | yes / yes |
| 250 | 299.6 | $D_t$ = 20% | 20.00 | 20.00 | 0.00 | 184.2 | 0.0 | no / no |
| 290 | 35.0 | operating point | 4.84 | 4.81 | −0.03 | 1.064 | −5.7 | yes / yes |
| 290 | 249.4 | $D_t$ = 20% | 19.95 | 20.01 | +0.06 | 250.5 | +0.8 | yes / no |
| 330 | 26.6 | operating point | 4.61 | 4.61 | 0.00 | 1.024 | −2.1 | yes / yes |
| 330 | 213.8 | $D_t$ = 20% | 20.03 | 20.00 | −0.03 | 329.8 | +1.0 | no / no |

Error, surrogate minus FEM; pp, percentage points. 150 and 290 µm are not in the training grid.
